# N-terminally truncated helper NLR *NRG1C* antagonizes immunity mediated by its full-length neighbors *NRG1A* and *NRG1B*

**DOI:** 10.1101/2021.01.27.428547

**Authors:** Zhongshou Wu, Lei Tian, Xin Li

## Abstract

Both animals and plants utilize nucleotide-binding leucine-rich repeat immune receptors (NLRs) to perceive the presence of pathogen-derived molecules and induce immune responses. *NLR* genes are far more abundant and diverse in higher plants. Interestingly, truncated NLRs, which lack one or more of the canonical domains, are also commonly encoded in plant genomes. However, little is known about their functions, especially regarding the N-terminally truncated ones. Here, we show that *Arabidopsis thaliana* (*A. thaliana*) N-terminally truncated helper NLR gene *NRG1C* (*N REQUIREMENT GENE 1*) is highly induced upon pathogen infection and in autoimmune mutants. The immune response and cell death conferred by some TIR (Toll/interleukin-1 receptor)-type NLRs (TNLs) are compromised in the *NRG1C* overexpression lines. Detailed genetic analysis revealed that NRG1C antagonizes the immunity mediated by its full-length neighbors *NRG1A* and *NRG1B*. Biochemical tests indicate that NRG1C possibly interferes with the EDS1-SAG101 complex, which likely signals together with NRG1A/1B. Interestingly, Brassicaceae NRG1Cs are functionally exchangeable, and the *Nicotiana benthamiana* (*N. benthamiana*) N-terminally truncated helper NLR NRG2 antagonizes NRG1 in tobacco. Together, our study uncovers an unexpected negative role of N-terminally truncated helper NLRs in different plants.

## Introduction

Plants utilize a sophisticated innate immune system to defend against a plethora of pathogens. They depend on immune receptors to recognize non-self and initiate defense. Plasma membrane localized pattern-recognition receptors (PRRs) can perceive pathogen-associated molecular patterns (PAMPs), leading to PAMP/pattern-triggered immunity (PTI) (Couto and Zipfel, 2016). On the other hand, intracellular nucleotide-binding leucine-rich repeat immune receptors (NLRs) are responsible for warding off adapted pathogens by detecting their secreted effectors, thereby activating effector-triggered immunity (ETI) (Cui et al., 2015).

NLRs are present in both animals and plants. They are members of the STAND (signal transduction ATPases with numerous domains) family consisting of a variable N-terminal domain, a central nucleotide-binding (NB) motif and a C-terminal leucine-rich repeat (LRR) region (Bonardi et al., 2012; Jones and Dangl, 2006). Typical plant sensor NLRs (sNLRs) have either a Toll/interleukin-1 Receptor (TIR) or a coiled-coil (CC) domain at their N-termini, and are termed TNLs (TIR-NB-LRR) or CNLs (CC-NB-LRR), respectively (Jones et al., 2016). They have evolved to recognize effectors directly or indirectly (Baggs et al., 2017). In contrast, a small subclass of NLRs is genetically required for diverse sNLRs’ immune signaling. These NLRs are termed helper NLRs (hNLR) (Bonardi et al., 2011; Collier et al., 2011; Jubic et al., 2019; Peart et al., 2005).

There are two *hNLR* gene families in *A. thaliana*, encoding RNLs (RPW8 (Resistance to Powdery Mildew 8) CC-NB-LRR): the *ACTIVATED DISEASE RESISTANCE 1* (*ADR1*) family (Bonardi et al., 2011) and the *N REQUIREMENGT GENE 1* (*NRG1*) family (Peart et al., 2005). Besides RNLs, the Solanaceae-specific CNL NRC (NB-LRR protein required for HR-associated cell death) proteins also function as hNLRs since both cell surface PRRs (Rathjen and Dodds, 2017; Wu et al., 2016) and many sNLRs signal through NRCs (Gabriëls et al., 2006; Gabriëls et al., 2007; Wu et al., 2017a). In comparison to the highly divergent sensor TNLs and CNLs, hNLRs are evolutionarily more conserved (Adachi et al., 2019; Andolfo et al., 2019; Shao et al., 2016; Wu et al., 2017a; Zhong and Cheng, 2016).

Plant *sNLRs* are among the most rapidly evolving genes (Gao et al., 2018; Guo et al., 2011; Van de Weyer et al., 2019). As a result, they are highly polymorphic. *NLR* allelic diversity mainly depends on the generation of either chimeric LRRs by intra- and intergenic recombination and gene conversion (Serra et al., 2018), or novel LRRs through point mutations (Lu et al., 2016; Mago et al., 2015; Michelmore and Meyers, 1998; Tamborski and Krasileva, 2020). Gene clustering is another feature commonly observed for plant *NLRs* (Van de Weyer et al., 2019). Two types of cluster patterns exist based on the diversity of *NLRs* within (Jacob et al., 2013). Homogenous clusters contain the same type of *NLRs* derived from tandem duplications. In contrast, heterogeneous clusters contain diverse *NLRs* generated by ectopic or large-scale segmental duplications (Jacob et al., 2013; Krasileva, 2019; Marone et al., 2013). Pan-genome analysis reveals that *NLR* cluster patterns vary among *A. thaliana* accessions. Some clusters have highly conserved copy numbers, while others show huge variations (Lee and Chae, 2020). An increasing number of examples suggest that clustered NLRs often are co-expressed and may cooperate to function. Clustering *sNLRs* together has the potential to massively expand pathogen recognition by increasing their oligomerization options (van Wersch and Li, 2019).

Duplication and uneven crossover events may generate *NLRs* lacking one or more of the canonical domains, yielding truncated *NLRs* (Baggs et al., 2017; Wicker et al., 2007), which are commonly found in all plant genomes (Bai et al., 2002; Jacob et al., 2013; Lee and Chae, 2020; Meyers et al., 2003; Meyers et al., 2002; Van de Weyer et al., 2019; Yang et al., 2008). Interesting, most truncated NLRs are expressed (Meyers et al., 2002; Nandety et al., 2013; Tan et al., 2007), suggesting their potential role in plant immunity. Several C-terminally truncated NLRs have been studied. For instance, *A. thaliana* TIR-only protein RBA1 (RESPONSE TO HOPBA1) is specifically required for defense responses mediated by bacterial effector HopBA1 (Nishimura et al., 2017). Another well-studied example includes the *A. thaliana* TN (TIR-NB) proteins CHS1 (CHILLING SENSITIVE 1) and TN2. CHS1 and TN2 form distinct pairs with full-length TNL SOC3 (SUPPRESSOR of *chs1-2*, 3) to monitor the homeostasis of E3 ligase SAUL1 (SENESCENCE-ASSOCIATED E3 UBIQUITIN LIAGASE 1) (Liang et al., 2019; Tong et al., 2017). However, little is known about the functions of N-terminally truncated NLRs, including NL (NB-LRR) and LRR proteins.

Although hNLRs may not have effector recognition capabilities, the NRG1-type *hNLRs* are organized in clusters in various plant species (Andolfo et al., 2019; Jubic et al., 2019; van Wersch and Li, 2019; Wu et al., 2019). In contrast, *ADR1s* are scattered in the genome (Bonardi et al., 2011; Dong et al., 2016). There are three *NRG1* paralogs in the *A. thaliana* genome: full length *NRG1A* and *NRG1B,* and N-terminally truncated *NRG1C* (Jubic et al., 2019; Wu et al., 2019). *NRG1A* and *NRG1B* are present in the *NRG1* cluster across almost all analyzed *A. thaliana* and *A. lyrate* accessions (Lee and Chae, 2020).

*NRC1C* encodes a predicted protein missing the RPW8 CC domain and most of the NB domain. Similarly, *N. benthamiana* contains a copy of the full-length *NRG1* and an N-terminally truncated *NRG2*. In comparison to the clustered *NRG1s* in *A. thaliana*, *NRG2* does not cluster with *NRG1* and has been assumed to be a pseudogene (Peart et al., 2005). It is unclear whether these N-terminally truncated hNLRs have any biological roles.

Here, we describe the functional analysis of truncated hNLRs. To our surprise, NRG1C negatively regulates TNL autoimmune mutant *chs3-2D* (*chilling sensitive 3, 2D*)-mediated autoimmunity, serving an opposite role from its full-length paralogs NRG1A/1B. *NRG1C* overexpression plants phenocopy those of the *nrg1a nrg1b* double mutant or the *sag101* (*SENESCENCE-ASSOCIATED GENE 101*) single mutant.

Biochemical analysis indicates that NRG1C possibly works by interfering with the EDS1 (ENHANCED DISEASE SUSCEPTIBILITY 1)-SAG101 complex. Interestingly, Brassicaceae NRG1Cs from different species are functionally exchangeable. In addition, the *N. benthamiana* truncated NRG2 also antagonizes NRG1-mediated cell death. In summary, our study uncovers an unexpected role of N-terminally truncated hNLRs, which was likely evolved to balance the activities of their corresponding full-length hNLRs.

## Results

### *NRG1C* is evolutionarily conserved in Brassicaceae and its expression is highly induced in TNL-type autoimmune mutants and upon pathogen infection

BLASTp analysis revealed that NRG1C (AT5G66890)-type N-terminally truncated hNLRs are present in all 8 well-sequenced Brassicaceae species and *N. benthamiana* (Figure 1A-B and Supplemental Figure 1). Additionally, *NRG1C* is present in most *A. thaliana* accessions. However, no NRG1C ortholog was found in available protein databases of other land plants. Interestingly, as with *A. thaliana NRG1C*, other Brassicaceae *NRG1C* orthologs also tend to cluster with the full length *NRG1* genes in their genomes (Supplemental Table 1). Phylogenetic analysis indicates that all Brassicaceae NRG1Cs came from the same ancestor (Figure 1B). The Brassicaceae *NRG1C* genes likely originated from either an initial gene duplication from the *NRG1A/1B* ancestor and a subsequent N-terminal loss, or partial gene duplication. In contrast, NbNRG2 and NbNRG1 belong to a distinct clade from Brassicaceae NRG1, suggestive of an independent evolution of NbNRG1 (Figure 1B and Supplemental Figure 1). In *Brassica rapa*, there are two close NRG1C homologs found in two separate gene clusters. Brara.I00877.1 and Brara.G01211.1 share 75.4% and 62.0% amino acid identity, or 86.2% and 75.5% similarity with NRG1C, respectively (Supplemental Figure 1). However, phylogenetic analysis indicates that Brara.G01211.1 might not be an NRG1C ortholog as it contains a full-length NB domain (Supplemental Figure 1) and is evolutionarily separate from the NRG1C clade (Figure 1B).

**Figure 1.**
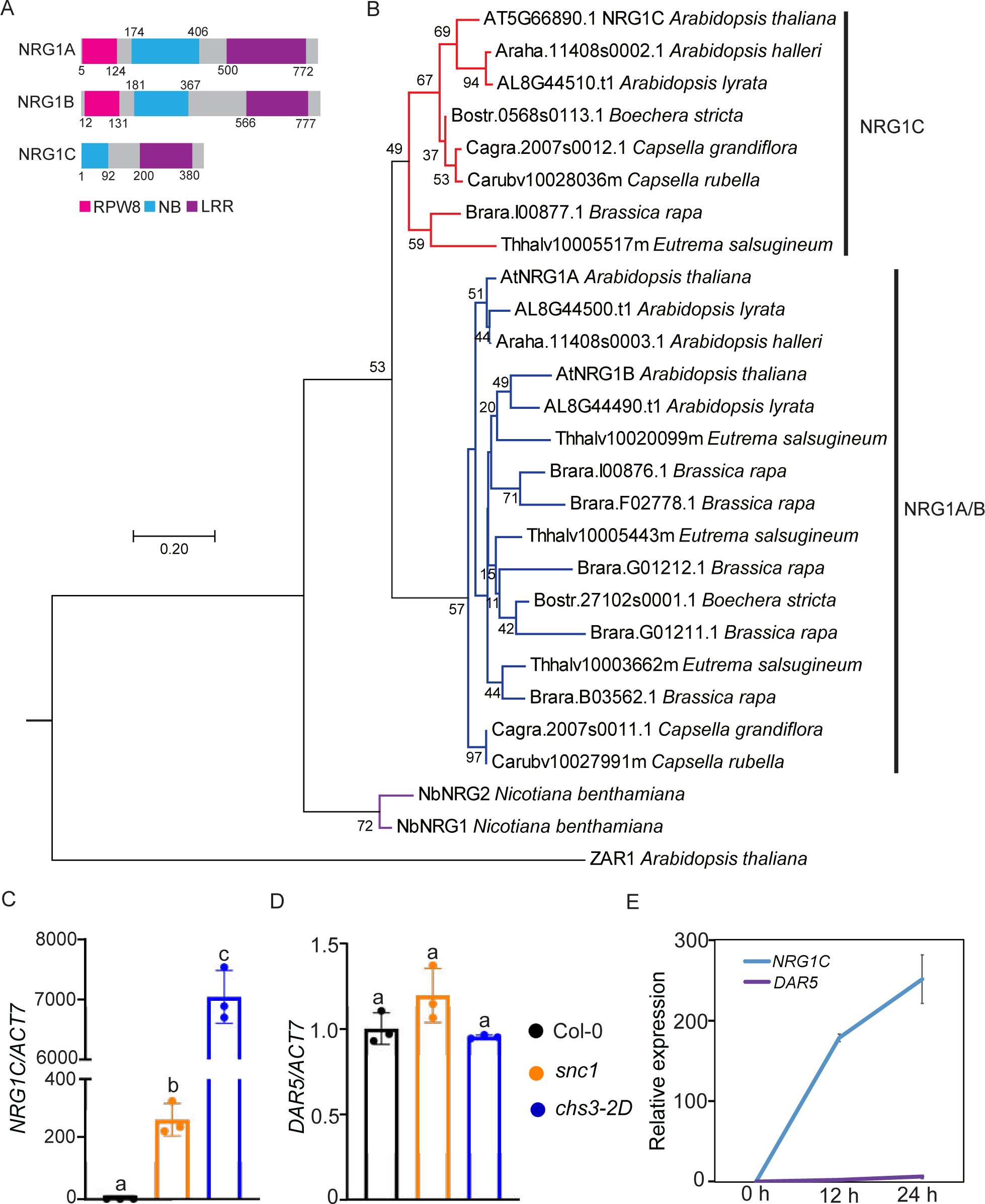
*NRG1C* is evolutionarily conserved in Brassicaceae, and its transcription is induced in TNL type autoimmune mutants and upon pathogen infection. (A) Protein domain diagram of NRG1A, NRG1B and NRG1C. Numbers represent amino acid positions relative to the translation start sites. NRG1C lacks the RPW8 CC and part of the NB domains. (B) Phylogenetic tree of putative NRG1A/1B/1C orthologs. Putative NRG1A/1B/1C orthologs were obtained from Phytozome (Brassicaceae species) and Sol Genomics Network (*N. benthamiana*) using *A. thaliana* NRG1A, NRG1B and NRG1C protein sequences as input, respectively. ZAR1 was used as an outgroup to root the tree. MUSCLE was used for sequence alignment and a Neighbor-joining tree was generated by using full-length NRG1C and homologous parts of all other corresponding NRG1 homologs with JTT model and using 2000 bootstrap replicates in MEGA 7.0. C-D. *NRG1C* (C) and *DAR5* (D) gene expression in the autoimmune mutants *snc1* and *chs3-2D* as determined by RT-PCR (Col-0 serves as wild type control and the transcript level in Col-0 was arbitrarily set at 1.0). Statistical significance is indicated by different letters (*p* < 0.01). Error bars represent means ± SE (Stand Error) (n = 3). Three independent experiments were carried out with similar results. (E) RT-PCR analysis of the expression of *NRG1C* and *DAR5* after *P.s.m.* ES4326 infection. Samples were collected at 0, 12 and 24 h after infection. Values were normalized to the level of *ACTIN7* and the transcript levels at 0 h post-inoculation (hpi) were set at 1.0. Error bars represent means ± SE (n = 3). Three independent experiments were carried out with similar results.

DAR5 (AT5G66630, DA1-RELATED PROTEIN 5) was proposed to be a chimeric protein resulting from a translocation of NRG1C and a fusion with a member of the nearby cluster carrying DAR homologs, DAR7 (AT5G66610) to DAR3 (AT5G66640) (Meyers et al., 2003). Coincidently, DAR5 carries RPW8, NB and LIM (Lin11, Isl-1 & Mec-3) domains, but no LRRs. It shares high similarity with NRG1s in *A. thaliana* (Andolfo et al., 2019; Castel et al., 2019). However, DAR5 seems to be unique in *A. thaliana*, as BLAST searches failed to identify putative DAR5 orthologs in genomes of all other sequenced plant species.

To test whether the truncated *NRG1C* and *DAR5* are involved in TNL-mediated immunity, their expression levels were first measured in TNL autoimmune mutants. *snc1* (*suppressor of npr1-1, constitutive 1*) and *chs3-2D* contain gain-of-function mutations in the *TNLs SNC1* and *CHS3*, respectively, resulting in TNL-mediated autoimmunity.

*NRG1C* transcripts were around 200 and 7000-fold up-regulated respectively in *snc1* and *chs3-2D* mutants as compared with that of Col-0 wild-type (WT) (Figure 1C). However, no significant *DAR5* transcript changes were observed (Figure 1D). Furthermore, the transcript of *NRG1C*, but not *DAR5*, was highly induced in WT plants upon infection by virulent bacterial pathogen *Pseudomonas syringae* pv. *maculicola* (*P.s.m.*) ES4326 (Figure 1E). These expression analyses suggest that *NRG1C* in the *NRG1* gene cluster may play roles in plant biotic responses against pathogens.

### *NRG1C*, but not *DAR5*, negatively regulates *chs3-2D* and *snc1*-mediated autoimmunity

As *NRG1C* is up-regulated in autoimmune mutants and during infection, and the *nrg1c* knockout does not seem to exhibit severe immune related defects and affect *snc1* autoimmunity (Wu et al., 2019), we used an overexpression approach to examine the role of these truncated NLRs. *NRG1C* and *DAR5* were individually overexpressed in *chs3-2D* and *snc1* backgrounds. To our surprise, overexpression of *NRG1C* fully suppressed *chs3-2D*-mediated dwarfism (Figure 2A-C) and enhanced disease resistance to the oomycete pathogen *Hyaloperonospora arabidopsidis* (*H.a.*) Noco2 (Figure 2D). In contrast, no suppression of *chs3-2D*-mediated growth defects was observed upon *DAR5* overexpression (Supplemental Figure 2A-B). Furthermore, overexpressing *NRG1C* (Figure 2E) could partially suppress *snc1*-mediated dwarfism to a similar level as the *NRG1* deletion mutant (Wu et al., 2019) (Figure 2F-G). Consistently, enhanced disease resistance to *H.a.* Noco2 (Figure 2H) and *P.s.t.* DC3000 (Figure 2I) in *snc1* were partially suppressed upon *NRG1C* overexpression. However, the *snc1 DAR5 OE* lines did not show significant differences in growth compared with *snc1* (Supplemental Figure 2C-D). Taken together, NRG1C negatively regulates *chs3-2D* and *snc1*-mediated autoimmunity with different strengths. Intriguingly, these phenotypes closely resemble those of the *nrg1a nrg1b* double mutant and the *sag101* single mutant (Wu et al., 2019; Xu et al., 2015).

**Figure 2.**
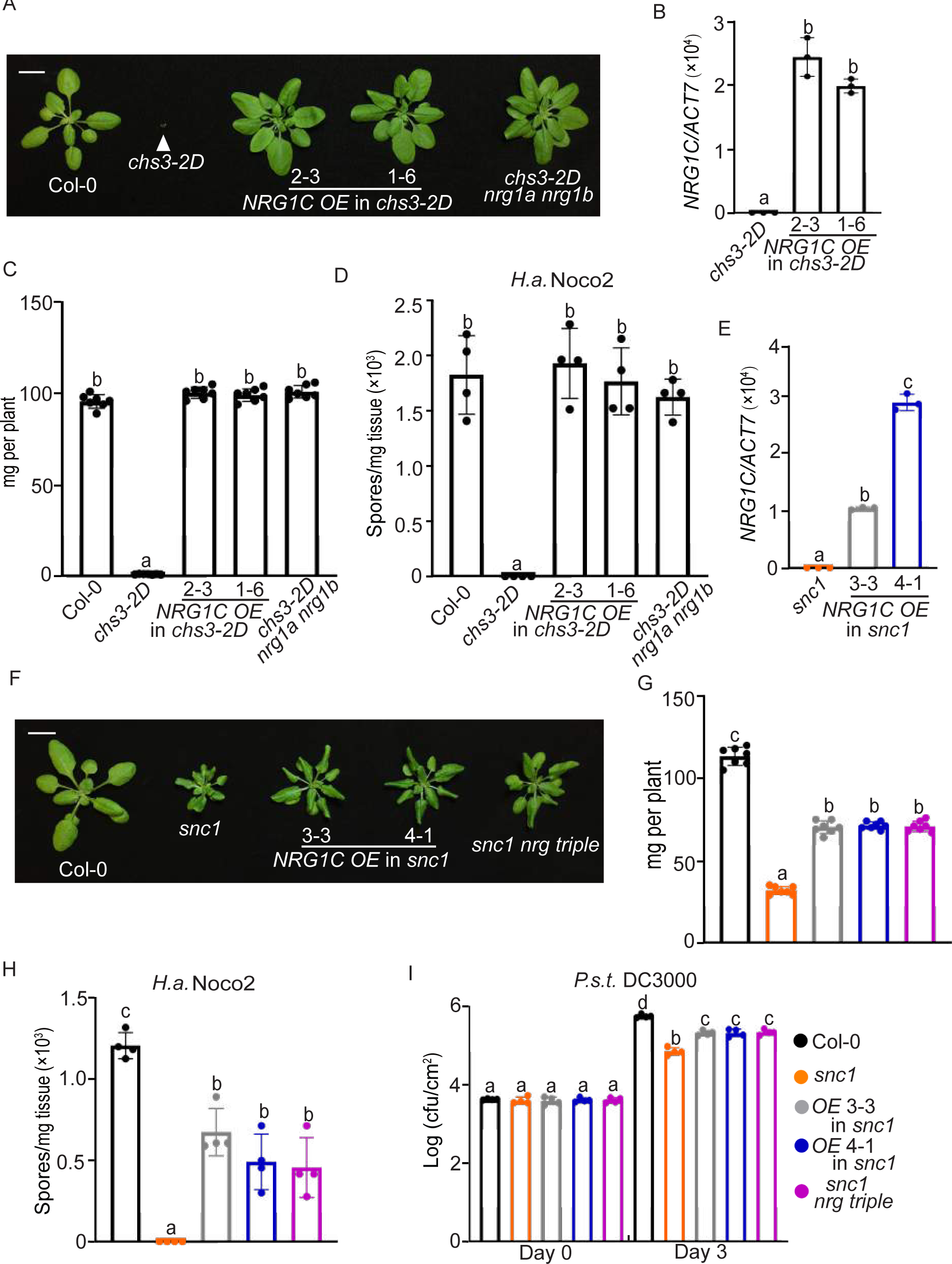
*NRG1C* negatively regulates *chs3-2D* and *snc1*-mediated autoimmunity. (A) Morphology of four-week-old soil-grown plants of Col-0, *chs3-2D*, two independent transgenic lines of *NRG1C OE* into *chs3-2D* background and *chs3-2D nrg1a nrg1b*. *OE* stands for overexpression of *NRG1C*. Bar = 1cm. (B) *NRG1C* gene expression in the indicated genotypes as determined by RT-PCR (*chs3-2D* serves as control whose *NRG1C* transcript level was set at 1.0). Statistical significance is indicated by different letters (*p* < 0.01). Error bars represent means ± SE (n = 3). Two independent experiments were carried out with similar results. (C) Fresh weights of plants in (A). Statistical significance is indicated by different letters (*p* < 0.01). Error bars represent means ± SE (n=7). (D) Quantification of *H.a.* Noco2 sporulation in the indicated genotypes 7 days post inoculation (dpi) with 10^5^ spores per ml water. Statistical significance is indicated by different letters (*p* < 0.01). Error bars represent means ± SE (n=4). Three independent experiments were carried out with similar results. (E) *NRG1C* gene expression in the indicated plants as determined by RT-PCR (*snc1* serves as control whose *NRG1C* transcript level was set at 1.0). Statistical significance is indicated by different letters (*p* < 0.01). Error bars represent means ± SE (n = 3). Two independent experiments were carried out with similar results. (F) Morphology of four-week-old soil-grown plants of Col-0, *snc1*, two independent transgenic lines of *NRG1C OE* into *snc1* background and *snc1 nrg triple*. Bar = 1cm. (G) Fresh weights of plants in (F). Statistical significance is indicated by different letters (*p* < 0.01). Error bars represent means ± SE (n=7). The color legends are included in I. (H) Quantification of *H.a.* Noco2 sporulation in the indicated genotypes at 7 dpi with 10^5^ spores per ml water. Statistical significance is indicated by different letters (*p* < 0.01). Error bars represent means ± SE (n=4). Three independent experiments were carried out with similar results. The color legends are included in I. (I) Bacterial growth of *P.s.t.* DC3000 in four-week-old leaves of the indicated genotypes at 0 and 3 dpi with bacterial inoculum of OD600 = 0.001. Statistical significance is indicated by different letters (*p* < 0.01). Error bars represent means ± SE (n=4). Three independent experiments were carried out with similar results.

### ADR1-L3 is dispensable for *snc1*-mediated autoimmunity and RPP4-mediated immunity

Similar to NRG1C in the NRG1 family, the *A. thaliana* ADR1 helper NLR family includes a truncated ADR1-L3, missing the N-terminal RPW8 domain (Supplemental Figure 3). In contrast to the broad presence of NRG1C in Brassicaceae, an ADR1-L3 ortholog is only found in *Boechera stricta* (Supplemental Figure 3). To test whether ADR1-L3 is involved in immunity conferred by ADR1-dependent NLRs, *ADR1-L3* was overexpressed in *snc1* background.

As shown in Supplemental Figure 4A-B, *snc1*-mediated growth defects (Supplemental Figure 4A) and enhanced disease resistance (Supplemental Figure 4B) were not altered upon *ADR1-L3* overexpression.

Furthermore, *ADR1-L3* overexpression lines in WT were not susceptible to oomycete pathogen *H.a.* Emwa1 (Supplemental Figure 4C), which is recognized by ADR1-dependent TNL RPP4 (*RECOGNITION OF PERONOSPORA PARASITICA 4*) (Bonardi et al., 2011; Dong et al., 2016). Taken together, unlike NRG1C, ADR1-L3 does not seem to play any roles in TNL-mediated autoimmunity and defense.

### Overexpression of *NRG1C* has no effect on *chs1-2*- and *chs2-1*-mediated autoimmunity, RPS2 and RPS4-mediated immunity, or basal defense

To further test the specificity of NRG1C, *NRG1C* was overexpressed in additional autoimmune mutants. In *chs1-2* (*chilling sensitive 1*, *2*), a missense mutation in a truncated TN protein (Wang et al., 2013) results in constitutive cell death and defense responses at low temperatures (Zbierzak et al., 2013). Another chilling sensitive mutant *chs2-1* contains a gain-of-function mutation in the TNL *RPP4* (Huang et al., 2010).

However, neither *chs1-2* nor *chs2-1*-mediated temperature-dependent autoimmunity was suppressed upon *NRG1C* overexpression (Supplemental Figure 5A-B). Consistently, oomycete pathogen *H.a.* Emwa1 failed to grow in *NRG1C OE* lines in the WT background (Supplemental Figure 5C-D).

We also challenged two independent *NRG1C OE* lines (in WT Col-0 background) with *P.s.t.* DC3000 expressing either AvrRps4 or AvrRpt2 effectors, which are recognized by TNL RPS4 (RESISTANT TO *P. SYRINGAE* 4) or CNL RPS2 (RESISTANT TO *P*. *SYRINGAE* 2), respectively. However, no altered disease resistance against these avirulent pathogens was detected (Supplemental Figure 6A-B). When challenged with *P.s.t.* DC3000, *NRG1C OE* lines exhibited similar bacterial growth as WT, suggesting that overexpression of *NRG1C* does not compromise basal defense response either (Supplemental Figure 6C). Therefore, NRG1C seems to be specific to TNL-mediated immune response with unequal strengths for distinct TNLs. The differential contributions of NRG1C in TNL signaling are reminiscent of the phenotypes of *nrg1a nrg1b* or *sag101* knockouts, which play major roles in *chs3-2D* but relatively mild roles in *snc1*, but no roles in *chs1-2*, *chs2-1*, RPS4 and RPS2-mediated autoimmunity and bacterial growth (Castel et al., 2019; Feys et al., 2005; Wu et al., 2019; Xu et al., 2015).

### NRG1C is required for HopQ1-1-mediated defense

NbNRG1 is essential for immune signaling triggered by effector HopQ1-1 (Qi et al., 2018). As *P.s.t.* DC3000 carries diverse effectors that can mask the effects of HopQ1-1, *P.s.t.* CUCPB5500 in which all 18 of the well-expressed T3SS (Type III secretion system) effectors were deleted, was used as a background to avoid interference from other effectors (Kvitko et al., 2009). When challenged with *P.s.t.* CUCPB5500, no bacterial growth difference was observed between *NRG1C OE* lines, *nrg1a nrg1b* double or *sag101-1* mutants as compared with WT (Figure 3A). However, WT exhibited enhanced disease resistance to *P.s.t.* CUCPB5500 expressing HopQ1-1, suggesting that unidentified R proteins can likely recognize HopQ1-1 in *A. thaliana*. Consistently with the observation in tobacco, this bacterial growth inhibition was largely compromised in *NRG1C OE* lines, *nrg1a nrg1b* double mutant and *sag101* single mutant to a similar level (Figure 3A). Taken together, NRG1C antagonizes HopQ1-1-mediated defense responses in *A. thaliana*. *NRG1C* overexpression phenotypes seem identical to those of the *nrg1a nrg1b* double and the *sag101* single mutants.

**Figure 3.**
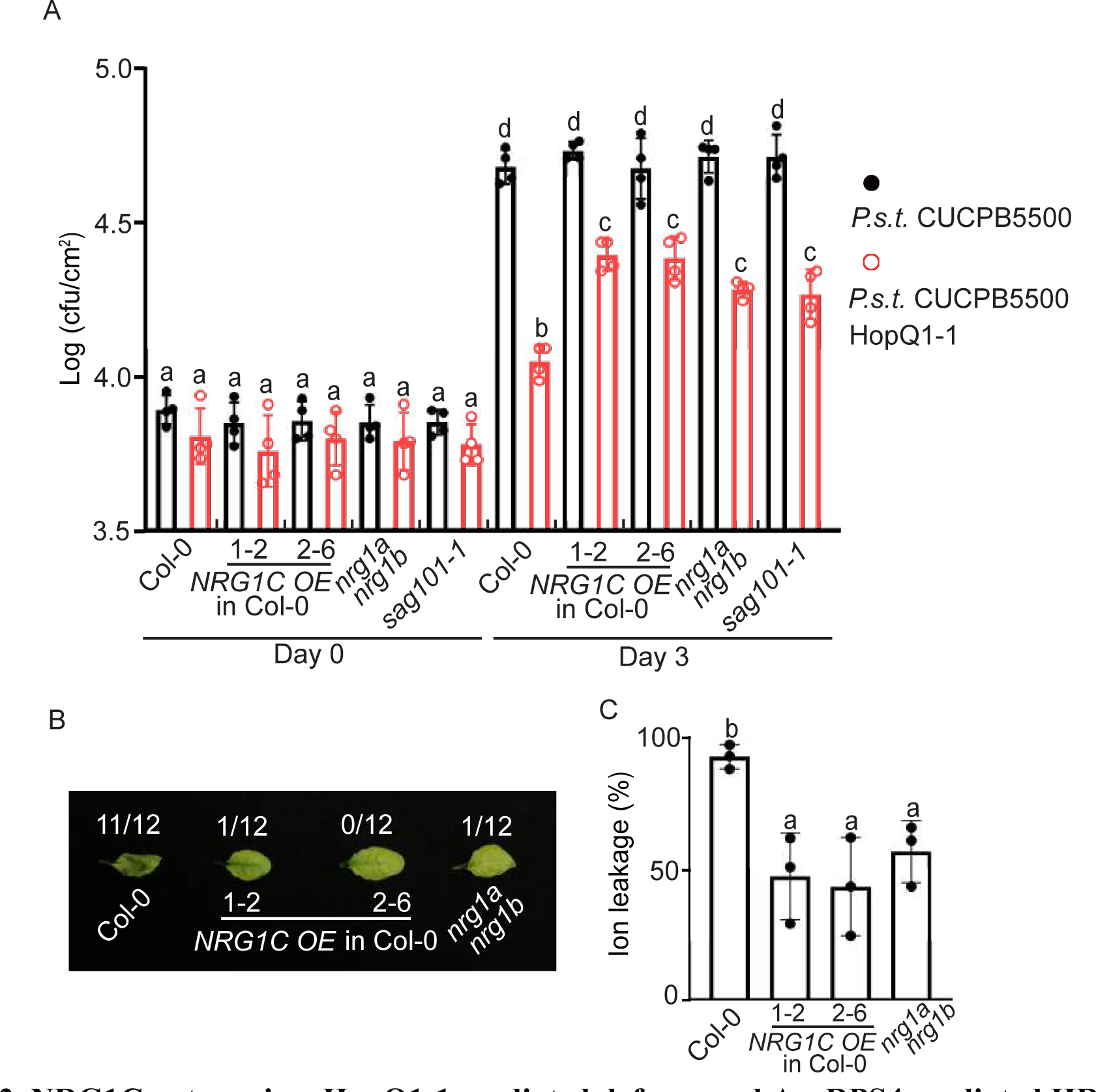
NRG1C antagonizes HopQ1-1-mediated defense and AvrRPS4-mediated HR. (A) Growth of *P.s.t.* CUCPB5500 and *P.s.t.* CUCPB5500 HopQ1-1 in four-week-old leaves of the indicated genotypes at 0 and 3 dpi with bacterial inoculum of OD600 = 0.0005. Statistical significance is indicated by different letters (*p* < 0.01). Error bars represent means ± SE (n=4). Three independent experiments were carried out with similar results. (B) HR observation of leaves infiltrated with *Pf0-1*_AvrRPS4. Four-week-old *A. thaliana* leaves of the indicated genotypes were infiltrated with *Pf0-1* expressing AvrRPS4 at OD600 = 0.2. Photographs were taken at 36 hpi. The numbers indicate the number of plants displaying HR out of the total number of plants tested. Three independent experiments were carried out with similar results. (C) Ion leakage of *A. thaliana* leaves upon *Pf0-1*_AvrRPS4 infiltration under the same conditions as for B. Each dot represents four technical replicates from one independent experiment. Statistical significance is indicated by different letters (*p* < 0.01). Error bars represent means ± SE (n=3). Three independent experiments were carried out with similar results.

### NRG1C antagonizes AvrRPS4-mediated hypersensitive response

AvrRPS4 can activate RRS1 (RESISTANT TO RALSTONIA SOLANACEARUM1)/RPS4 pairs, resulting in hypersensitive response (HR) in Col-0 background (Saucet et al., 2015). This HR is compromised in Col-0 *nrg1a nrg1b* double mutant, although no difference in *P.s.t.* AvrRPS4 bacterial growth can be observed (Supplemental Figure 6A) (Castel et al., 2019; Lapin et al., 2019). We thus tested whether NRG1C regulates RRS1/RPS4-mediated HR. Effector AvrRPS4 was delivered using *P. fluorescens* strain *Pf0-1* containing type III secretion system (*Pf0-1* EtHAn), but lacking type III effector genes (Thomas et al., 2009). Similar with the phenotype of *nrg1a nrg1b* double mutant (Castel et al., 2019; Lapin et al., 2019), RRS1/RPS4-mediated HR is lost in *NRG1C OE* lines (Figure 3B-C). The striking overall phenotypic resemblance between *NRG1C OE* lines and the *nrg1a nrg1b* double mutant indicates that NRG1C might function antagonistically with NRG1A/1B in *A. thaliana* to regulate defense responses.

### NRG1C antagonizes the NRG1-SAG101 module, in parallel with the ADR1-PAD4 module

The similar phenotypes of *adr1 triple* with *pad4* (Phytoalexin deficient 4) mutants (Dong et al., 2016), and the additive effects between SAG101 and PAD4 (Rietz et al., 2011) or NRG1s and ADR1s (Lapin et al., 2019; Saile et al., 2020; Wu et al., 2019) allude to an EDS1-ADR1-PAD4 signaling module that works in parallel with an EDS1-NRG1-SAG101 module to regulate downstream events in TNL immunity. To test this further, higher order mutants combining loss-of-function mutations from different modules were generated in *snc1* background. The dwarfism and increased *PR1* (*Pathogenesis-Related 1*) gene expression of *snc1* was partially suppressed by introducing mutations in *SAG101* (Feys et al., 2005) or *NRG1* cluster (Wu et al., 2019) (Figure 4A-B). However, *snc1 sag101-1 nrg triple* resembled the *snc1 sag101-1* and *snc1 nrg triple* in morphology (Figure 4A) and *PR1* gene expression levels (Figure 4B), confirming that SAG101 works in the same pathway as NRG1A/1B. In contrast, the increased *PR1* gene expression of *snc1* was largely, but not fully, suppressed by *adr1 triple* or *pad4-1* (Figure 4B), although the size of *snc1 adr1 triple* and *snc1 pad4-1* was WT like (Cheng et al., 2011; Dong et al., 2016) (Figure 4A). However, combining *adr1 triple* and *pad4* mutations together did not further reduce the *PR1* gene expression (Figure 4B and S7), confirming that ADR1s and PAD4 work in the same pathway. In contrast, mutants containing mutations in two components from different modules, including *snc1 pad4-c1 sag101-1*, *snc1 pad4-c1 nrg triple*, *snc1 sag101-c1 adr1 triple*, *snc1 adr1 triple nrg triple*, exhibited fully restored WT-like levels of *PR1* expression (Figure 4A-B and S7). Taken together, these epistasis analyses data confirm that two distinct modules are present in TNL pathways that act in parallel, one includes PAD4-ADR1s and the other includes SAG101-NRG1A/1B.

**Figure 4.**
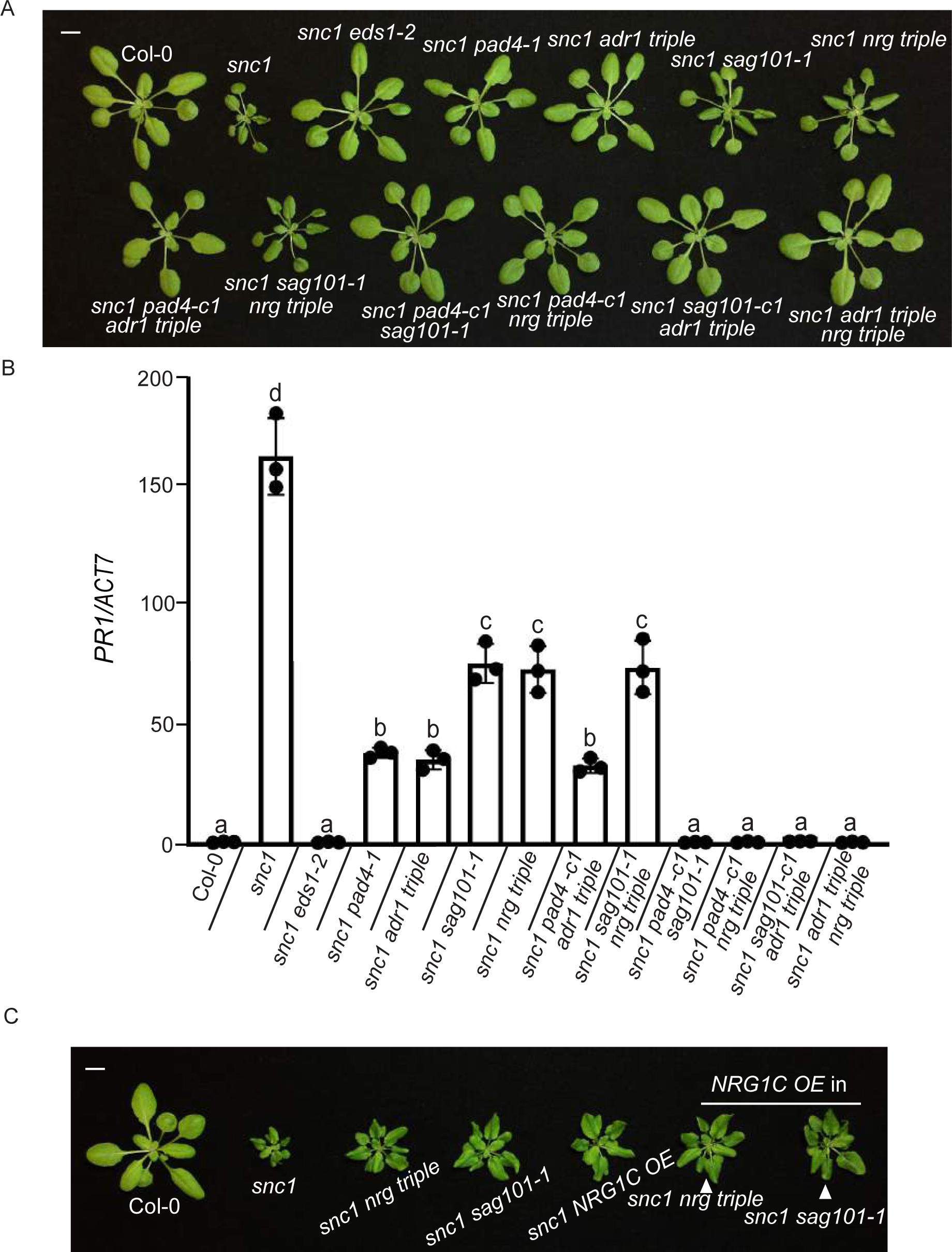
NRG1C works along the EDS1-NRG1-SAG101 module, in parallel with the EDS1-ADR1-PAD4 module. (A) Morphology of four-week-old soil-grown plants of Col-0, *snc1*, *snc1 eds1-*2, *snc1 pad4-1*, *snc1 adr1 triple*, *snc1 sag101-1*, *snc1 nrg triple*, *snc1 pad4-c1 adr1 triple*, *snc1 sag101-1 nrg triple*, *snc1 pad4-c1 sag101-1*, *snc1 pad4-c1 nrg triple*, *snc1 sag101-c1 adr1 triple* and *snc1 adr1 triple nrg triple*. *pad4-c1* and *sag101-c1* alleles were generated through the CRISPR-Cas9 system. Bar = 1cm. (B) *PR1* gene expression in the indicated genotypes as determined by RT-PCR (Col-0 serves as control whose *PR1* transcript level was set at 1.0). Statistical significance is indicated by different letters (*p* < 0.01). Error bars represent means ± SE (n = 3). Two independent experiments were carried out with similar results. (C) Morphology of four-week-old soil-grown plants of Col-0, *snc1*, *snc1 nrg triple*, *snc1 sag101-1*, *snc1 NRG1C* OE, *snc1 nrg triple NRG1C OE* and *snc1 sag101-1 NRG1C OE*. *OE* stands for overexpression of *NRG1C*. Bar = 1cm.

To examine the relationship between NRG1C and the NRG1-SAG101 module, *NRG1C* was overexpressed in the *snc1 nrg triple* or *snc1 sag101-1* background. As shown in Figure 4C, the dwarfism was not further suppressed by *NRG1C* overexpression. In agreement, *snc1 sag101-1 NRG1C OE* and *snc1 nrg triple NRG1C OE* resembled the *snc1 sag101-1* and *snc1 nrg triple* in regards to *PR1* gene expression (Supplemental Figure 8A). Specifically, the increased *PR1* gene expression in *snc1* was reduced by either *nrg triple* or *sag101*, but not further by *NRG1C* overexpression (Supplemental Figure 8A). In contrast, the partial suppression of up-regulated *PR1* gene expression in *snc1* by *adr1 triple* or *pad4-1* was further suppressed to WT levels when *NRG1C* was overexpressed in *snc1 adr1 triple* or *snc1 pad4-1* backgrounds (Supplemental Figure 8B). These comprehensive epistasis data support that NRG1C indeed specifically antagonizes the NRG1-SAG101 module.

### *NRG1C* does not impact *NRG1A/1B* through transcriptional interference

As *NRG1C* encodes a truncated hNLR and works antagonistically with the SAG101- NRG1 module, we first hypothesized that it might interfere with *NRG1A/1B* transcription through RNA-mediated interference due to their sequence similarity. To test this, *NRG1C* with a premature stop codon at position 213 of the predicted *NRG1C* mRNA sequence was overexpressed in *chs3-2D* background. Such a mutation would yield a non-functional protein NRG1C^T71*^ but a rather normal transcript. No alterations in *chs3-2D*-mediated dwarfism (Figure 5A-C) and enhanced disease resistance to *H.a.* Noco2 (Figure 5D) were observed by overexpression of *NRG1C^T71*^* (Figure 5B), suggesting that NRG1C does not affect mRNA levels of the full-length helper *NRG1A/1B*. Thus, if NRG1C interferes with NRG1A/1B, the effects are likely protein-related.

**Figure 5.**
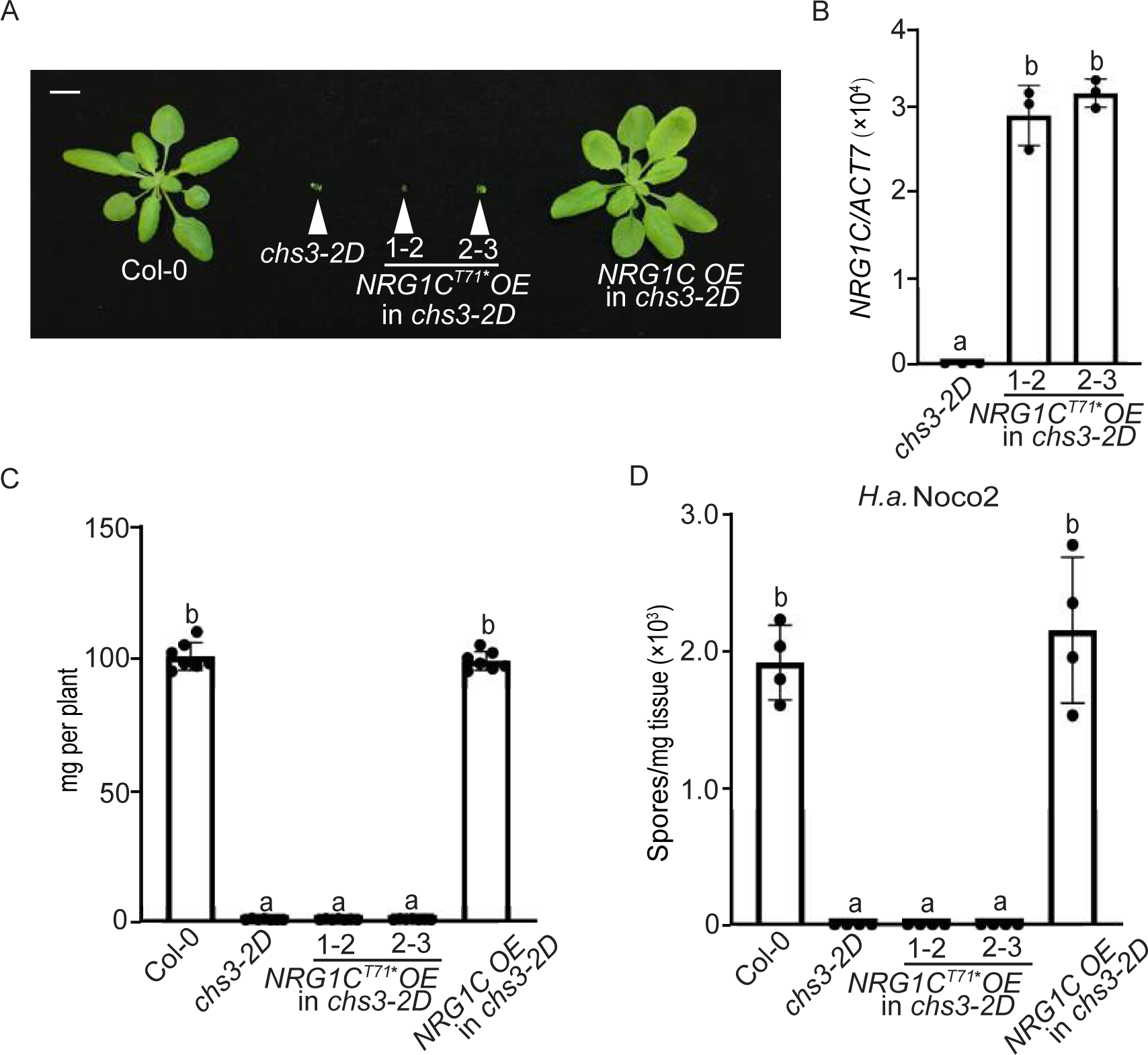
NRG1C regulates *chs3-2D*-mediated autoimmunity through processes unrelated to transcriptional interference. (A) Morphology of four-week-old soil-grown plants of Col-0, *chs3-2D*, and two independent transgenic lines of *NRG1C^T71*^ OE* into *chs3-2D* background. *OE* stands for overexpression of *NRG1C^T71*^*, which carries an early stop codon to disrupt protein synthesis, but not transcription. Bar = 1cm. (B) *NRG1C^T71*^* gene expression in the indicated genotypes as determined by RT-PCR (*chs3-2D* serves as control whose *NRG1C* transcript level was set at 1.0). Statistical significance is indicated by different letters (*p* < 0.01). Error bars represent means ± SE (n = 3). Two independent experiments were carried out with similar results. (C) Fresh weights of plants in (A). Statistical significance is indicated by different letters (*p* < 0.01). Error bars represent means ± SE (n=7). (D) Quantification of *H.a.* Noco2 sporulation in the indicated genotypes at 7 dpi with 10^5^ spores per ml water. Statistical significance is indicated by different letters (*p* < 0.01). Error bars represent means ± SE (n=4). Three independent experiments were carried out with similar results.

### NRG1C antagonizes NRG1A-mediated defense and HR

To further examine the hypothesis that NRG1C might directly interfere with NRG1A/1B, we overexpressed *NRG1C* in autoimmune transgenic plants expressing *NRG1A D485V*, an auto-active version of NRG1A that constitutively activates immunity (Wu et al., 2019). The dwarfism and enhanced resistance of *NRG1A D485V* were largely suppressed upon *NRG1C* overexpression (Figure 6A-B and S9A), while no effect on NRG1A protein level was observed (Figure 6C). These results further support that the negative regulation of NRG1A by NRG1C is not through transcriptional or post-transcriptional repression.

**Figure 6.**
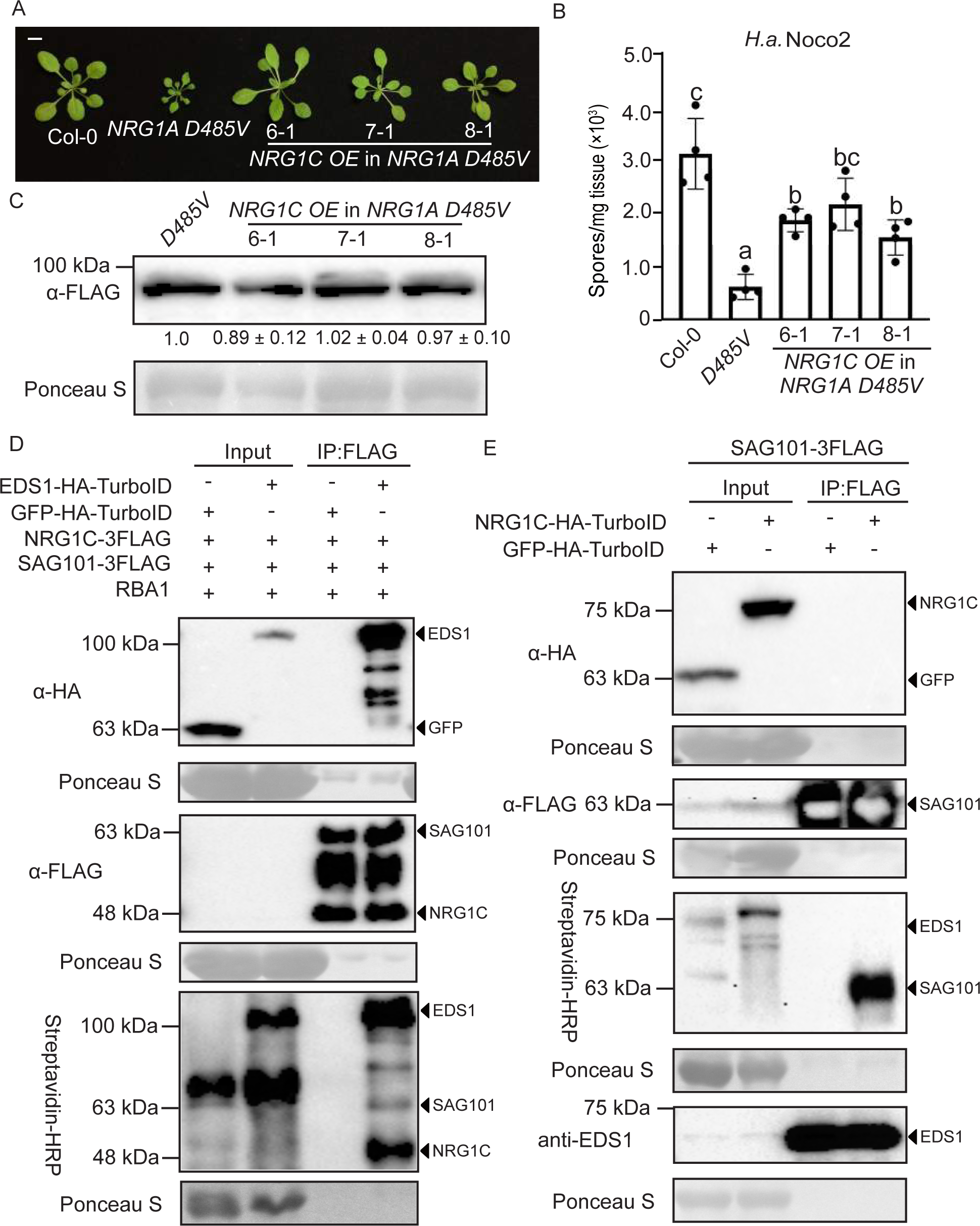
NRG1C antagonizes NRG1A *D485V*-mediated autoimmunity through associating with EDS1-SAG101 complex. (A) Morphology of three-week-old soil-grown plants of Col-0, *NRG1A D485V*, and three independent transgenic lines of *NRG1C OE* into *NRG1A D485V* background. *OE* stands for overexpression of *NRG1C*. *NRG1A D485V* was tagged with three FLAG epitopes in frame at its C-terminus. Bar = 1cm. (B) Quantification of *H.a.* Noco2 sporulation in the indicated genotypes at 7 dpi with 10^5^ spores per ml water. Statistical significance is indicated by different letters (*p* < 0.01). Error bars represent means ± SE (n=4). Three independent experiments were carried out with similar results. (C) Immunoblot analysis of NRG1A D485V protein levels in the indicated genotypes. Equal loading is shown by Ponceau S staining of a non-specific band. The numbers below represent the normalized ratio between the intensity of the protein band and the Ponceau S band ± SE (n=3). Molecular mass marker in kiloDaltons is indicated on the left. (D) Immunoprecipitation and biotinylation of NRG1C-3FLAG and SAG101-3FLAG by EDS1-HA-TurboID in *N. benthamiana* in the presence of RBA1. Immunoprecipitation was carried out with anti-FLAG beads. The 3FLAG-tagged proteins were detected using an anti-FLAG antibody. The HA-TurboID-tagged proteins were detected using an anti-HA antibody. The biotinylated proteins were detected using Streptavidin-HRP. Molecular mass marker in kiloDaltons is indicated on the left. The experiment was repeated three times with similar results. (E) Biotinylation of SAG101-3FLAG by NRG1C-HA-TurboID in *A. thaliana* plants stably transformed with the two transgenes in the presence of RBA1. Immunoprecipitation was carried out with anti-FLAG beads. The 3FLAG-tagged proteins were detected using an anti-FLAG antibody. The HA-TurboID-tagged proteins were detected using an anti-HA antibody. EDS1 was detected using an anti-EDS1 antibody. The biotinylated proteins were detected using HRP-Streptavidin. Molecular mass marker in kiloDaltons is indicated on the left. The experiment was repeated twice with similar results.

Consistent with our observation in *A. thaliana*, NRG1A D485V-triggered HR in *N. benthamiana* was alleviated when co-expressed with NRG1C (Supplemental Figure 9B-C), although the same amount of NRG1A D485V proteins accumulated (Supplemental Figure 9D). However, overexpression of *NRG1C* did not alter the autoimmunity of *ADR1-L2 D484V*, an auto-active variant of ADR1-L2 (Roberts et al., 2013) (Supplemental Figure 10A-C). Taken together, these data indicate that NRG1A-mediated autoimmunity and HR are negatively regulated by NRG1C and the effects of NRG1C seem to be specific to NRG1-type hNLR-mediated defense responses.

### NRG1C does not associate with full-length NRG1A

Since NRG1C does not affect the mRNA or protein levels of full-length NRG1A, another hypothesis is that NRG1C might interfere with NRG1A/1B through protein-protein interactions. The recently developed TurboID-based proximity labeling method allows TurboID-fused protein to biotinylate proximal and interacting proteins in the presence of biotin *in vivo* (Zhang et al., 2019). This unbiased method can more reliably detect weak and transient protein-protein interactions. When NRG1A-HA-TurboID was co-expressed with NRG1C-3FLAG in the presence of RBA1, which induces NRG1 dependent immunity (Figure S11A) (Qi et al., 2018; Wan et al., 2019), biotinylated NRG1C-3FLAG was not detected despite the fact that NRG1C-3FLAG protein was significantly enriched (Supplemental Figure 11B). This negative interaction result suggests that the NRG1C-mediated negative regulation of NRG1A-mediated defense may not be through direct protein interference. However, we cannot exclude the possibility that the *in planta* interactions between the two proteins could not be detected due to low protein expression or complex conformation constraints.

### NRG1C associates with the EDS1-SAG101 complex

As *NRG1C* overexpression also shares similar phenotypes with the *sag101* knockouts, we then tested whether NRG1C could interfere with NRG1A-mediated defense by associating with SAG101 or its constitutively associated partner EDS1. When *N. benthamiana* leaves were infiltrated with *Agrobacterium* expressing EDS1-HA-TurboID, NRG1C-3FLAG and SAG101-3FLAG, both IP-enriched SAG101-3FLAG and NRG1C-3FLAG were biotinylated (Supplemental Figure 12A). This suggests that basal levels of NRG1C proteins are in close proximity to the C-terminus of EDS1, where the TurboID was fused. In comparison, in *N. benthamiana* leaves pre-infiltrated with *Agrobacterium* harboring RBA1, a similar amount of IP-enriched SAG101-3FLAG, but relatively more NRG1C-3FLAG, was biotinylated by EDS1-HA-TurboID (Figure 6D, Supplemental Figure 12A and Supplemental Figure 14C), suggesting that TNL activation likely enhances the association between NRG1C and the EDS1-SAG101 complex. Interestingly, when this TurboID-mediated proximity labeling assay was performed in the absence of SAG101-3FLAG, biotinylated NRG1C was no longer detected (Supplemental Figure 12B), indicating that the association between EDS1 and SAG101 is needed to retain NRG1C in the complex.

We further examined such interaction with a reciprocal TurboID-based proximity labeling assay using *A. thaliana* stable transgenic lines carrying both *NRG1C-HA-TurboID* and *SAG101-3FLAG*. Transgenic plants were pretreated with *Agrobacterium* expressing RBA1 and whole seedling plants were harvested at 6 hpi. As shown in Figure 6E, although NRG1C-HA-TurboID could not be observed in the SAG101-3FLAG pull-down likely due to its low protein abundance, NRG1C-HA-TurboID successfully biotinylated SAG101-3FLAG, but not EDS1, even though a large amount of EDS1 protein was pulled down. This suggests that SAG101 is in close proximity with the C-terminus of NRG1C, where TurboID is fused. In addition, neither EDS1 (Supplemental Figure 13A-B) nor SAG101 (Supplemental Figure 13C-D) protein levels were altered by *NRG1C* overexpression, confirming that NRG1C does not affect the protein levels of the EDS1-SAG101-NRG1A/1B module.

TIR domains of many TNLs exhibit nicotinamide adenine dinucleotide (NAD^+^) hydrolase (NADase) activity, with a conserved catalytic glutamic acid (Horsefield et al., 2019; Wan et al., 2019). The catalytically dead RBA1E86A mutant retains self-association, but fails to trigger cell death *in planta* (Wan et al., 2019). To test whether the association between NRG1C and the EDS1-SAG101 complex is influenced by the TIR NADase activity, *N. benthamiana* leaves expressing EDS1-HA-TurboID, SAG101-3FLAG and NRG1C-3FLAG were pretreated with *Agrobacterium* expressing *RBA1E86A* mutant. As shown in Supplemental Figure 14A and S14C, SAG101-3FLAG and basal levels of NRG1C-3FLAG were biotinylated by EDS1-HA-TurboID, suggesting that TIR NADase enzyme activity is required for TNL-enhanced high order complex formation.

As EDS1 also constitutively interacts with PAD4, we also tested whether NRG1C associates with the EDS1-PAD4 complex. However, when *N. benthamiana* leaves were infiltrated with *Agrobacterium* expressing EDS1-HA-TurboID, NRG1C-3FLAG and PAD4-ZZ-TEV-FLAG in the presence of RBA1 induction, EDS1-HA-TurboID was able to label PAD4-ZZ-TEV-FLAG, but not NRG1C-3FLAG (Supplemental Figure 14B). This specificity of NRG1C with SAG101 explains why NRG1C explicitly antagonizes EDS1-SAG101-NRG1 module-dependent immunity. It is worth noting that *N. benthamiana* contains its own copies of *NRG1*, *SAG101* and *EDS1*. NbSAG101 is apparently not sufficient to induce proximity between AtEDS1 and AtNRG1C (Supplemental Figure 12B and S14B). This is consistent with previous findings that lipase-like proteins (EDS1, PAD4, and SAG101) and hNLRs (ADR1s and NRG1s) can only signal together when they come from the same species (Lapin et al., 2019).

### Overexpression of *B. rapa* NRG1C ortholog suppresses *chs3-2D*-mediated defense

As the presence of *NRG1C*-type N-terminally truncated hNLRs in *NRG1* clusters seems to be a singular event in the evolution of Brassicaceae, we further tested whether they are functionally transferable among different Brassicaceae species. To test this, *Brara.I00877.1* (*BraNRG1C*) was overexpressed in the *chs3-2D* mutant. Consistent with the phylogeny that Brassicaceae NRG1C family arose from a common ancestor (Figure 1A), overexpression of *BraNRG1C* fully rescued the *chs3-2D*-mediated dwarfism (Figure 7A-C) and enhanced disease resistance to *H.a.* Noco2 (Figure 7D). Taken together, NRG1C function seems to be conserved in Brassicaceae species.

**Figure 7.**
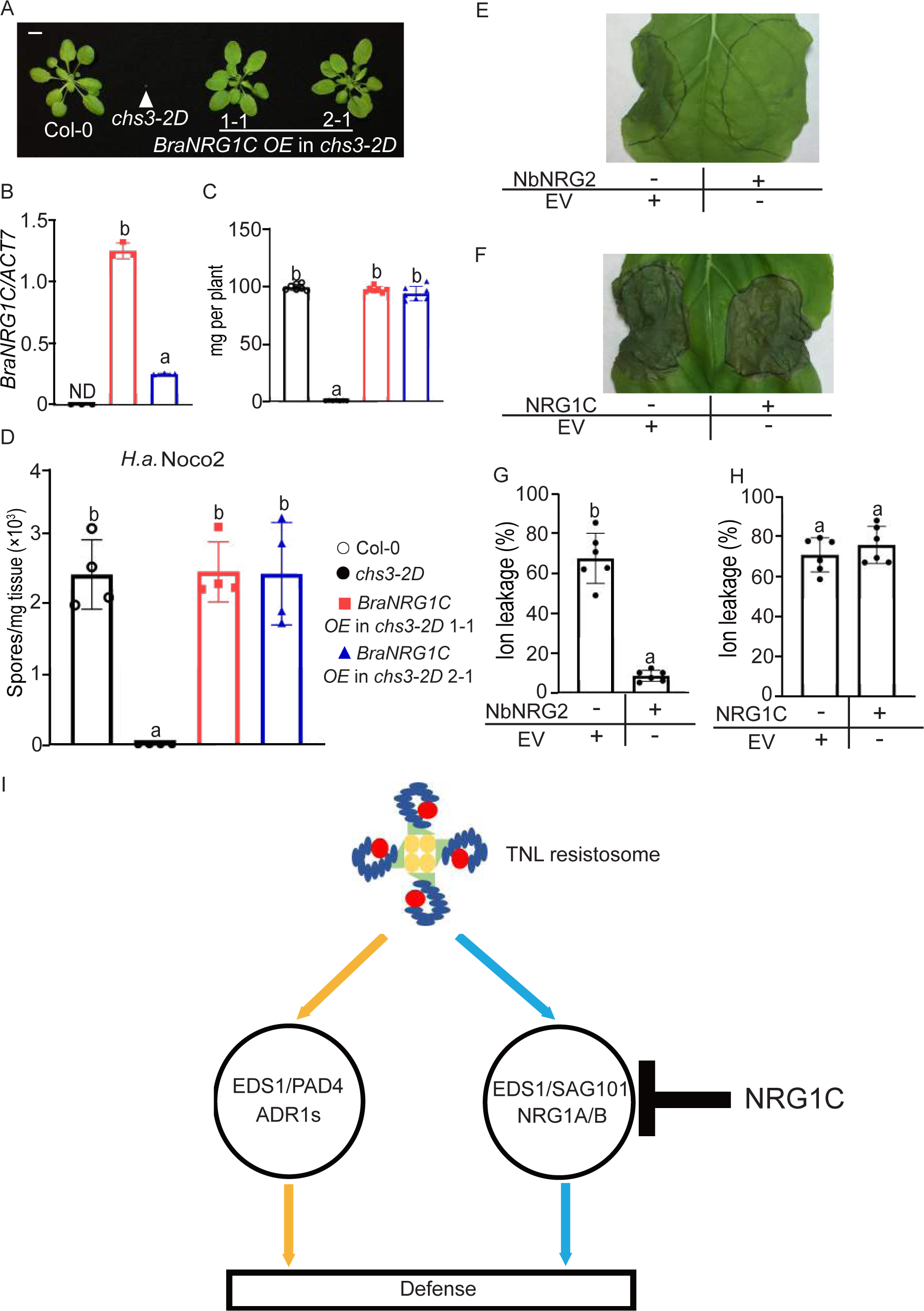
NRG1C-type N-terminally truncated hNLR from canola (*Brassica napus* L. cv. Westar) functions similarly as *A. thaliana* NRG1C. (A) Morphology of four-week-old soil-grown plants of Col-0, *chs3-2D*, and two independent transgenic lines of *BraNRG1C OE* into *chs3-2D* background. Bar = 1cm. (B) *BraNRG1C* gene expression in the indicated plants as determined by RT-PCR. Statistical significance is indicated by different letters (*p* < 0.01). Error bars represent means ± SE (n = 3). ND stands for Not Detectable. Two independent experiments were carried out with similar results. The color legends are included in D. (C) Fresh weights of plants in (A). Statistical significance is indicated by different letters (*p* < 0.01). Error bars represent means ± SE (n=7). The color legends are included in D. (D) Quantification of *H.a.* Noco2 sporulation in the indicated genotypes at 7 dpi with 10^5^ spores per ml water. Statistical significance is indicated by different letters (*p* < 0.01). Error bars represent means ± SE (n=4). Three independent experiments were carried out with similar results. E-F. HR observation of *N. benthamiana* leaves expressing the indicated proteins at 48 hpi following the infiltration of *P.s.t.* CUCPB5500 HopQ1-1 to deliver HopQ1-1 effector. Photographs were taken at 4 dpi after *P.s.t.* CUCPB5500 HopQ1-1 infiltration. G-H. Ion leakage of *N. benthamiana* leaves upon *P.s.t.* CUCPB5500 HopQ1-1 infiltration under the same conditions as with E and F. Statistical significance is indicated by different letters (*p* < 0.01). Error bars represent means ± SE (n = 6). Three independent experiments were carried out with similar results. (I) A model of two defense modules downstream of TNL activation in Arabidopsis. An effector-induced TNL resistosome with NADase activity (Horsefield et al., 2019; Ma et al., 2020; Martin et al., 2020; Tian and Li, 2020; Wan et al., 2019) signals through two parallel modules, EDS1-PAD4-ADR1s and EDS1-SAG101-NRG1A/1B to activate defense responses. N-terminally truncated hNLR NRG1C antagonizes immunity mediated by its full-length neighbors NRG1A and NRG1B to avoid aberrant activation.

### NbNRG2 negatively regulates HopQ1-1-mediated immunity in *N. benthamiana*

To further explore further the function of truncated NRG1C outside the Brassicaceae, we examined *N. benthamiana* NRG2 (NbNRG2) in HopQ1-1-mediated NRG1-dependent ROQ1 (RECOGNITION of XopQ 1) resistance (Qi et al., 2018; Schultink et al., 2017) (Supplemental Figure 15A). Interestingly, overexpression of *NbNRG2* (Supplemental Figure 15B) could suppress HopQ1-1-mediated HR, which was induced by *P.s.t.* CUCPB5500 HopQ1-1 (Figure 7E and G). By contrast, the HopQ1-1-mediated HR was not suppressed by overexpression of Brassicaceae orthologs including *A. thaliana NRG1C* (Figure 7F and H; Supplemental Figure 15C) or *BraNRG1C* (Supplemental Figure 15D-F), indicating that these Brassicaceae orthologs are not able to inhibit *N. benthamiana* TNL ROQ1-mediated immunity. Furthermore, overexpression of *NbNRG2* (Supplemental Figure 16B) failed to suppress *chs3-2D*-mediated dwarfism (Supplemental Figure 16A and C) and enhanced disease resistance to *H.a.* Noco2 (Supplemental Figure 16D). Since molecular incompatibility among different species or clades was observed in the EDS1-SAG101-NRG1 module (Lapin et al., 2019), it is not surprising that NbNRG2 and Brassicaceae NRG1Cs are not functionally exchangeable and their negative regulation requires molecular compatibility.

## Discussion

Truncated NLRs, including TIR only, TN, TIR-unknown domain (TX), CC only, CC-NB (CN), CC-unknown domain (CX), RPW8 only, RPW8-NB (RN), NB-LRR (NL), NB only and LRR only, are commonly found encoded in the genomes of higher plants (Baggs et al., 2017; Bai et al., 2002; Jacob et al., 2013; Lee and Chae, 2020; Meyers et al., 2003; Meyers et al., 2002; Nandety et al., 2013; Van de Weyer et al., 2019; Yang et al., 2008). A genome-wide survey of *A. thaliana* NLR genes revealed that around 31% of NLR genes are truncated (Meyers et al., 2003). Among them, approximately 90% lack canonical C-terminal domains (Meyers et al., 2003; Meyers et al., 2002). Functional characterizations of some C-terminally truncated NLRs have been reported. The TIR-only protein RBA1 can function as an effector sensor and trigger plant immune responses by self-oligomerization (Nishimura et al., 2017). Moreover, the TIR domain in the RBA1 protein possesses NADase activity, which is required to trigger cell death (Wan et al., 2019). Other well-studied TN proteins include TN2 and CHS1. TN2 guards the exocyst complex subunit EXO70B1 (Zhao et al., 2015) and may stabilize active calcium-dependent protein kinase CPK5 to regulate defense responses (Liu et al., 2017).

Interestingly, *TNL SOC3* and *CHS1* cluster together with *TN2* (Liang et al., 2019; Zhang et al., 2017). SOC3 pairs with either CHS1 or TN2 to monitor the absence or overaccumulation of E3 ligase SAUL1 (Liang et al., 2019; Tong et al., 2017). A recent study found that LRR-PL (Post-LRR) truncated protein DM10^TueScha-9^ (DANGEROUS MIX 10) shows an incompatible interaction with DM11^Cdm-0^ (DANGEROUS MIX 11), resulting in hybrid necrosis (Barragan et al., 2020). Therefore, the C-terminally truncated NLRs often can still serve positive roles in NLR activation, as they retain the N-terminal signaling domains such as the TIR and NB regions.

In contrast, about 10% of truncated NLRs are N-terminally truncated in *A. thaliana* (Meyers et al., 2003). Although they are widely present in all higher plant genomes, little is known about their biological significance. In mammalian system, the TLR (Toll-like receptors) TIR truncations can act as dominant-negative (DN) proteins to dampen immune responses by associating with the same immune activation complexes (Burns et al., 2003; Padmanabhan et al., 2009; Wells et al., 2006). Here, similar to the truncated TLRs, the N-terminally truncated helper NRG1C lacking the RPW8 CC and part of the NB domain can negatively regulate NRG1A/1B-mediated defense responses. Compared to the DN-TLR examples in the mammalian system, how exactly NRG1C interferes with NRG1A/1B signaling is less clear. As overexpression of a *NRG1C* variant containing a premature stop codon failed to suppress *chs3-2D*-mediated autoimmunity, the mechanism is unlikely to be due to transcriptional interference (Figure 5). Additionally, NRG1A-HA-TurboID was unable to label NRG1C-3FLAG even in the presence of an elicitor, suggesting that NRG1C does not interact with NRG1A/1B, although caution should always be taken with negative protein-protein interaction data. However, NRG1C is in close proximity with the EDS1-SAG101 complex, and this association seems to be enhanced with TNL-induction (Figure 6D-E, Supplemental Figure 12A-B and Supplemental Figure 14C). To our delight, our observation was recently independently supported by a SAG101-YFP IP-MS (Immunoprecipitation Mass-Spectrometry) analysis, where peptides of NRG1C were abundantly observed in immuno-precipitates of SAG101-YFP upon *Pf0-1 avrRPS4* treatment (Figure 3a and Supplemental Figure 5A, Sun et al., 2020). More peptides from NRG1C were present in the SAG101 IP than the ones from NRG1A/1B, corroborating the interaction between NRG1C and EDS1-SAG101. Additionally, Sun *et al*., also found that NRG1A associates with the EDS1-SAG101 dimer in an effector-dependent manner. Although EDS1 is required for all tested TNLs, the SAG101-NRG1 branch plays unequal roles in different cases (Feys et al., 2005; Wu et al., 2019; Xu et al., 2015) For example, SAG101 and NRG1A/1B are fully required for chs3-2D-mediated autoimmunity, but they are not needed for CHS1-SOC3 or RPP4-mediated autoimmunity in *saul1-1* or *chs2-1* (Wu et al., 2019). Therefore, the EDS1-SAG101 dimer is specifically required for some, but not all TNLs-mediated immunity and NRG1C likely antagonizes NRG1A/1B signaling by interfering with the EDS1-SAG101 dimer. It seems that upon infection, a large complex consisting of EDS1, NRG1A/1B and SAG101 forms, and NRG1C likely acts by interfering with such complex assembly to avoid aberrant activation (Figure 7I). NRG1C may occupy the NRG1A/1B space in the EDS1-SAG101-NRG1 complex, preventing NRG1A/1B interaction with the lipase-like proteins. This explains the negative interaction data between NRG1A and NRG1C. Further biochemical and structural analyses of these proteins are needed to clarify the details of these interactions.

Both NRG1C and ADR1-L3 are N-terminally truncated hNLRs. Although NRG1C negatively regulates some TNL-mediated defense, no effect on ADR1-dependent TNL mediated autoimmunity or pathogen growth was observed upon *ADR1-L3* overexpression (Supplemental Figure 4A-C). In contrast to NRG1C, ADR1-L3 retains the full NB domain (Supplemental Figure 1 and S3). NRG1C orthologs are commonly found in Brassicaceae (Figure 1A), whereas ADR1-L3 ortholog is only present in *Boechera stricta* (Supplemental Figure 3). Such an evolutionary difference indicates that ADR1-L3 is likely a pseudogene and thus was lost during evolution. Therefore, for N-terminally truncated NLRs in higher plant genomes, some may be pseudogenes, and some may have negative functions as with NRG1C. Although an evolutionary examination may provide an initial hint, functional analysis is needed to determine their exact roles.

Besides NRG1C and ADR1-L3, five additional N-terminally truncated *NB-LRR* genes are present in the *A. thaliana* Col-0 genome (Supplemental Table 2). AT4G19050, AT5G45510 and AT1G61300 are more closely related to CNLs (Meyers et al., 2003). *AT4G19050* is adjacent to a CN gene *AT4G19060*. *AT5G45510* and head-to-head arranged CNL *AT5G45520* form a cluster with two other CN genes *AT5G45490* and *AT5G45440*.

*AT1G61310* and adjacent CNL *AT1G61300*, are transcribed in the same direction (Supplemental Table 2). Only two NL proteins, AT5G38350 and AT4G09360, share a similar NB motif with TNLs. *AT5G38350* clusters together with TIR-only gene *AT5G38344* and TNL gene *AT5G38340*, while *AT4G09360* resides in the TNL cluster containing TNL *AT4G09430* and *AT4G09420* (Supplemental Table 2). Therefore, clustering of *NL* genes with full-length or C-terminally truncated sNLRs can also be detected in *A. thaliana*. It is possible that the*NL* genes in these clusters plays a similar role as *NRG1C* in the *NRG1* cluster, serving negative regulatory functions. Additional investigations into the functions of such *NL* genes in *A. thaliana* and other plants will be of great interest to widen our understanding of NLRs.

## Materials and Methods

### Plant growth conditions

*A. thaliana* and *N. benthamiana* plants were grown at 22°C under 16hr light/8hr dark regime, unless otherwise specified. Severe autoimmune mutants were grown at 28 °C to suppress the dwarfism and enable seed production.

### Construction of plasmids

Overexpression construct of *NRG1C* was made in *pCambia1305* vector. *NRG1C^T71*^* was cloned into *pCambia1305* vector as well. The overexpression construct of *NRG1C* in *NRG1A D485V* was cloned into *pBASTA* vector. The overexpression constructs of *BraNRG1C* and *NbNRG2* were cloned into *pBASTA* vector by using the extracted genomic DNA from *Brassica napus L.* cv*. Westar* and *N. benthamiana*. *DAR5* was cloned into *pHan* vector. For the TurboID-based proximity labeling assay, *NRG1A* and *EDS1* were cloned into *pBASTA-HA-TurboID*. *NRG1C* was cloned into *pCambia1305-3FLAG*. *pBASTA NRG1A-3FLAG* and *pCambia1300 EDS1-3FLAG* were described in previous studies (Wu et al., 2019).

To generate the CRISPR/Cas9 constructs for knocking out *PAD4* and *SAG101*, genomic sequences of *PAD4* and *SAG101* were subjected to CRISPRscan (http://www.crisprscan.org/?page=sequence) to identify the target sequences. The selected sequences were evaluated with Cas-OFFinder (http://www.rgenome.net/cas-offinder/). The pHEE401E vector was used to generate the CRISPR/Cas9 constructs following the protocols previously described (Wang et al., 2015). All primers used are listed in Supplemental Table 3.

### *A. thaliana* stable transformation

The binary constructs were introduced into *Agrobacterium tumefaciens* GV3101 by electroporation and subsequently transformed into *A. thaliana* by the floral dipping method (Clough and Bent, 1998). For *snc1*, *chs1-2* and *chs2-1* transformation, plants were grown at room temperature. For *chs3-2D* transformation, plants were grown at 28°C under 16hr light/8hr dark regime to reduce autoimmunity. Transformants were selected either on soil by spraying BASTA (Glufosinate ammonium) or on plates with Hygromycin B. 10 transformants were selected each and co-segregation analysis in T2 generations were carried out to make sure the suppression phenotypes were due to the transgene overexpression.

### Transient *N. benthamiana* expression

Transient *N. benthamiana* expression assays were performed as previously described (Wu et al., 2017b). In brief, *Agrobacteria* carrying the respective constructs were co-infiltrated into four-week-old *N. benthamiana* leaves. Final OD600 for most strains was 0.2, whereas OD600 =1.0 and 0.6 were used for all strains expressing NRG1C or SAG101, respectively. Each infiltration contained *Agrobacteria* expressing *p19* to inhibit gene silencing (Lakatos et al., 2004).

### Pathogen infections

Oomycete and bacterial pathogen infection assays were carried out as described previously (Li et al., 2001). In brief, two-week-old soil-grown seedlings were sprayed with *H.a.* Noco2 or Emwa1 conidia spores at a concentration of 10^5^ spores per ml water. Sporulation was quantified using a hemocytometer after plants were grown at 18°C for 7 days. For bacterial infections, four-week-old plants were infiltrated with bacterial solution at designated concentrations. Leaf discs were collected and ground on the day of infection (Day 0) and 3 days later (Day 3). Colony-forming units (cfu) were calculated after incubation on LB plates with appropriate antibiotic selection. For HR assay in *A. thaliana*, four-week-old soil-grown plants were hand infiltrated with *Pf0-1* carrying pBS46:AvrRPS4. Cell death was monitored at 36 hpi.

### Expression analysis

About 0.05 g of four-week-old soil-grown plant tissue was collected, and RNA was extracted using an RNA isolation kit (Bio Basic; Cat#BS82314). ProtoScript II reverse transcriptase (NEB; Cat#B0368) was used to generate cDNA. Real-time PCR was performed using a SYBR premix kit (TaKaRa, Cat#RR82LR). The primers used are listed in Supplemental Table 3.

### *Agrobacteria* infiltration following *P.s.t.* CUCPB5500 HopQ1-1 infection

*Agrobacteria* carrying the respective constructs were infiltrated into four-week-old *N. benthamiana* leaves. *P.s.t.* CUCPB5500 HopQ1-1 was then infiltrated in the indicated area 48 hrs after *Agrobacteria* infiltration. Ion leakage was measured at 4 days after *P.s.t.* CUCPB5500 HopQ1-1 infection.

### Ion leakage measurement in *N. benthamiana*

After agrobacteria infiltration and *P.s.t.* CUCPB5500 HopQ1-1 infection, leaf discs from *N. benthamiana* were collected and washed in 10 ml of milliQ water overnight at room temperature. Ion leakage was measured with a conductometer VWR E C METER Model 2052. All samples were then autoclaved to measure the total ion leakage.

### TurboID-based proximity labeling in *N. benthamiana*

TurboID-based proximity labeling assay was performed as described previously (Wu et al., 2020; Zhang et al., 2019). In brief, *N. benthamiana* leaves were infiltrated with agrobacteria containing *HA-TurboID* and *3FLAG* tagged constructs. At 48 hpi, agrobacteria expressing RBA1 or RBA1E86A constructs were infiltrated (Wan et al., 2019). At 52 hpi, biotin was infiltrated, and the plants were incubated at room temperature for 2 h to allow labeling. Leaves were harvested, followed by immunoprecipitation and western blot analysis.

### TurboID-based proximity labeling in *Arabidopsis*

Two-week-old plate-grown *Arabidopsis* transgenic lines carrying both *NRG1C-HA-TurboID* and *SAG101-3FLAG* were soaked with *Agrobacteria* expressing RBA1 (OD600=0.2). At 4 hpi, biotin was added in solution and the plants were incubated at room temperature for 2 hours for labeling. The plants were then harvested, followed by immunoprecipitation and western blot analysis.

### Protein extraction, immunoprecipitation, and western blot analysis

100 mg soil-grown *A. thaliana* plant leaves or *N. benthamiana* leaves were collected and extracted by extraction buffer (100 mM Tris-HCl pH 8.0, 0.2% SDS and 2% β-mercaptoethanol). Loading buffer was added to each protein sample and boiled for 5 min. Protein abundance was quantified using ImageJ (https://imagej.nih.gov/ij/).

Co-immunoprecipitation assay was performed as previously described (Wu et al., 2017b). In brief, *N. benthamiana* plants were infiltrated with agrobacteria. About 2.5 g *N*.

*benthamiana* leaves expressing the indicated proteins were harvested at 54 hpi and ground into powder with liquid nitrogen. Extraction buffer contains 25 mM Tris-HCl pH 7.5, 150 mM NaCl, 1mM EDTA, 0.15% Nonidet P-40, 10% Glycerol, 1 mM PMSF, 1× Protease Inhibitor Cocktail (Roche; Cat. #11873580001), and 10 mM DTT. The FLAG-tagged NRG1C and SAG101 proteins were immunoprecipitated using 20 µl M2 beads (Sigma; Cat. #A2220) for co-immunoprecipitation and biotinylation was detected by Streptavidin-HRP (Abcam Cat. # ab7403) The anti-HA antibody was from Roche (Cat. #11867423001). The anti-FLAG antibody was from Sigma (Cat. #F1804). The anti-EDS1 antibody was generously shared from Dr. Jane E. Parker (Feys et al., 2001).

### Statistical analysis

One-way ANOVA followed by Tukey’s post hoc test was performed. The Scheffé multiple comparison was applied for testing correction. Normality test for all data was done in SPSS. Statistical significance was indicated with different letters. *p* values and sample numbers (n) were detailed in figure legends.

## Acknowledgments

Drs. Jeff Dangl, Jane Parker, Brian Staskawicz, Pingtao Ding, Jonathan Jones, Yu Ti Cheng and Shenyang He are cordially thanked for generous sharing of *A. thaliana* seeds of mutants, anti-EDS1 antibody and *Pseudomonas Avr* strains. Dr. Marc T. Nishimura is thanked for sharing the RBA1 constructs. Dr. Brian J. Staskawicz is thanked for sharing the *nrg1-1 N. benthamiana* seeds. Mr. Kevin Ao and Dr. Paul Kapos are sincerely thanked for careful reading of the manuscript. This work was financially supported by CFI-JELF, the Natural Sciences and Engineering Research Council of Canada (NSERC) Discovery Program, NSERC-CREATE PRoTECT program, and the WD Cooper Memorial Fund from UBC.

## Author contributions

ZW, Data curation, Validation, Investigation, Methodology, Writing - original draft, Project administration; LT, Validation, Methodology; X Li, Conceptualization, Data curation, Formal analysis, Supervision, Funding acquisition, Writing—original draft, Project administration, Writing - revisions. All authors reviewed the manuscript.

---

**Supplemental Figure 1.**
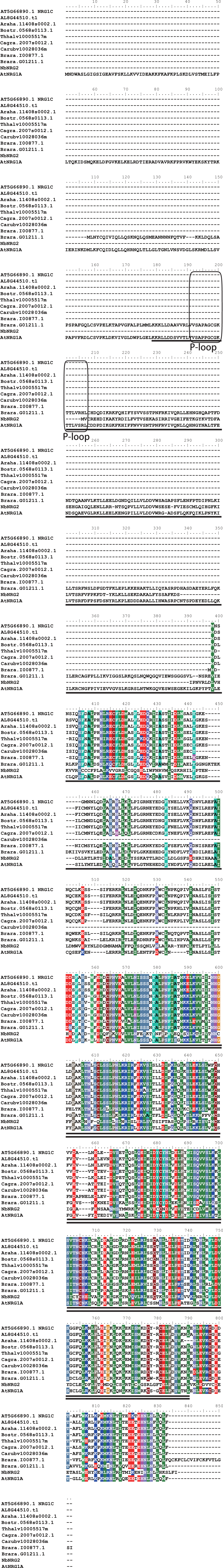
Full-length protein sequence alignment of NRG1C and its orthologs in different plant species. Putative NRG1C orthologs were obtained from Phytozome (Brassicaceae species) and Sol Genomics Network (*N. benthamiana*) using *A. thaliana* NRG1C protein sequence as input. NB domain and LRR domain are indicated by single underline and double underline, respectively. P-loop motif is boxed in a black line.

**Supplemental Figure 2.**
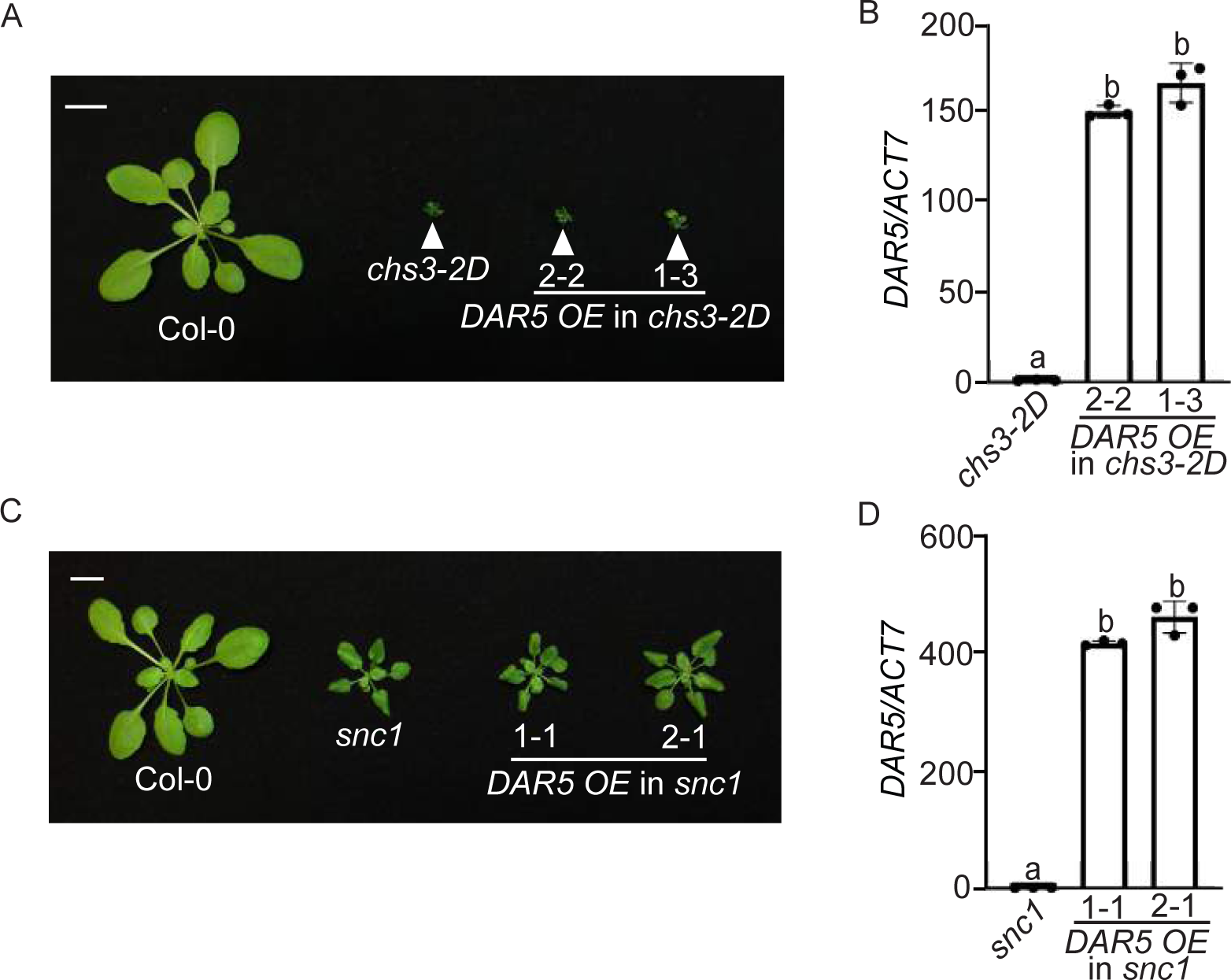
DAR5 has no effect on *chs3-2D* and *snc1*-mediated autoimmunity. (A) Morphology of four-week-old soil-grown plants of Col-0, *chs3-2D*, and two independent transgenic lines of *DAR5 OE* into *chs3-2D* background. *OE* stands for overexpression of *DAR5*. Bar = 1cm. (B) *DAR5* gene expression in the indicated plants as determined by RT-PCR (*chs3-2D* serves as control whose *DAR5* transcript level was set at 1.0). Statistical significance is indicated by different letters (*p* < 0.01). Error bars represent means ± SE (n = 3). Two independent experiments were carried out with similar results. (C) Morphology of four-week-old soil-grown plants of Col-0, *snc1*, and two independent transgenic lines of *DAR5 OE* into *snc1* background. *OE* stands for overexpression of *DAR5*. Bar = 1cm. (D) *DAR5* gene expression in the indicated plants as determined by RT-PCR (*snc1* serves as control whose *DAR5* transcript level was set at 1.0). Statistical significance is indicated by different letters (*p* < 0.01). Error bars represent means ± SE (n = 3). Two independent experiments were carried out with similar results.

**Supplemental Figure 3.**
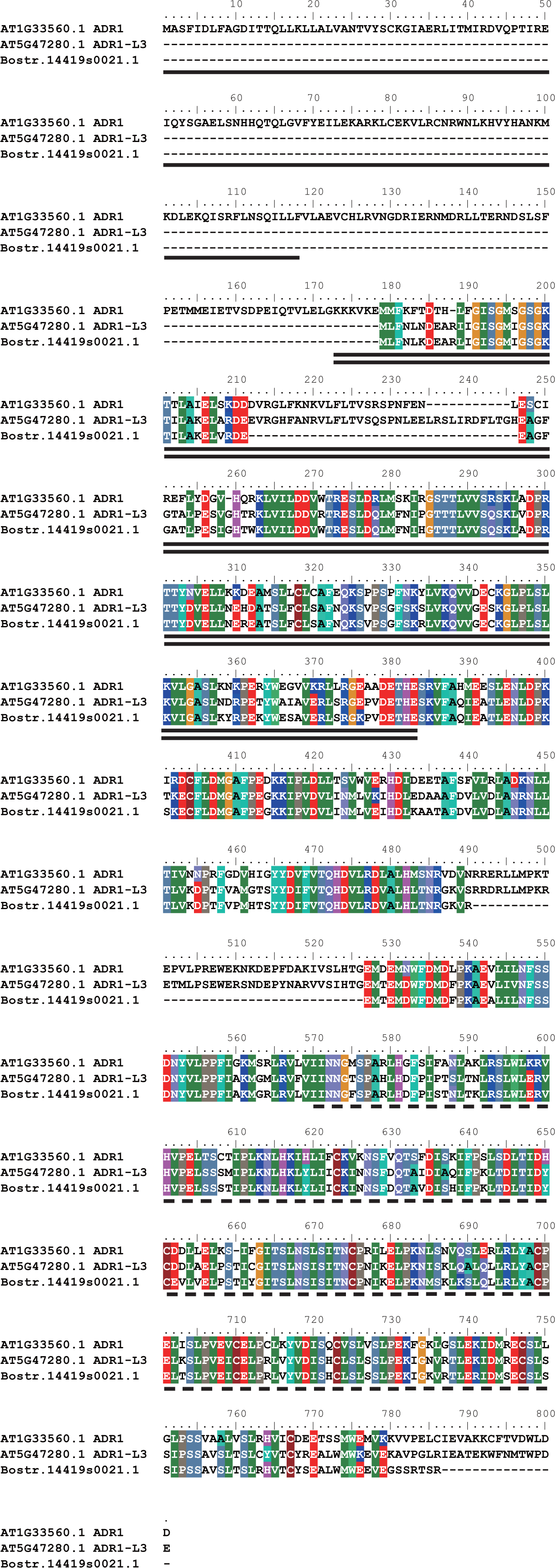
Full-length protein sequence alignment of ADR1-L3 and its orthologs in different plant species. Putative ADR1-L3 orthologs were obtained from Phytozome using *A. thaliana* ADR1-L3 protein sequence as input. RPW8 domain, NB domain and LRR domain are indicated by single underline, double underline and dashed underline, respectively.

**Supplemental Figure 4.**
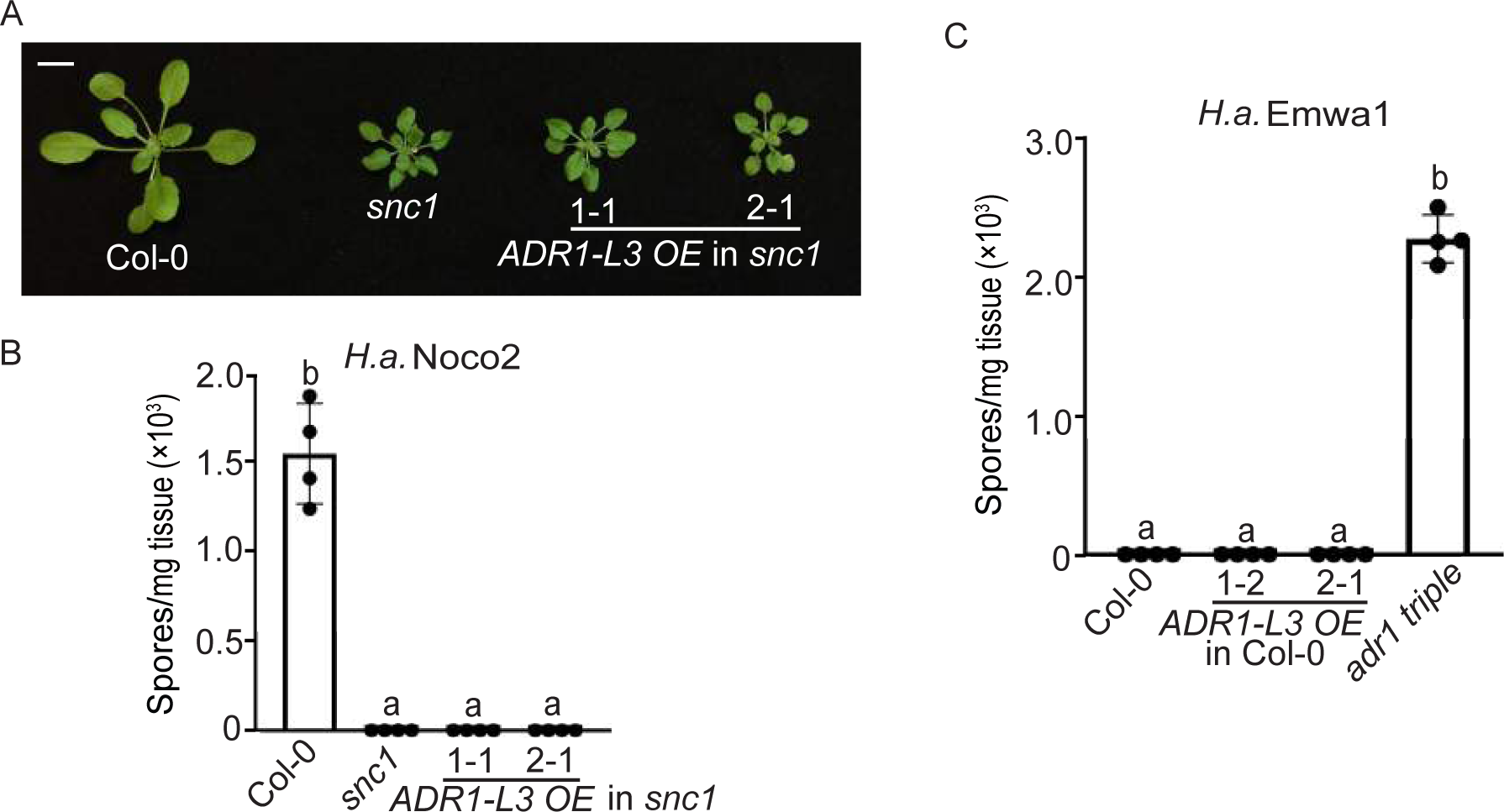
No immune-related phenotypes were observed in *ADR1-L3* overexpression lines. (A) Morphology of four-week-old soil-grown plants of Col-0, *snc1*, and two independent transgenic lines of *ADR1-L3 OE* into *snc1* background. *OE* stands for overexpression of *ADR1-L3*. Bar = 1cm. (B) Quantification of *H.a.* Noco2 sporulation in the indicated genotypes at 7 dpi with 10^5^ spores per ml water. Statistical significance is indicated by different letters (*p* < 0.01). Error bars represent means ± SE (n=4). Three independent experiments were carried out with similar results. (C) Quantification of *H.a.* Emwa1 sporulation in the indicated genotypes at 7 dpi with 10^5^ spores per ml water. Statistical significance is indicated by different letters (*p* < 0.01). Error bars represent means ± SE (n=4). Three independent experiments were carried out with similar results.

**Supplemental Figure 5.**
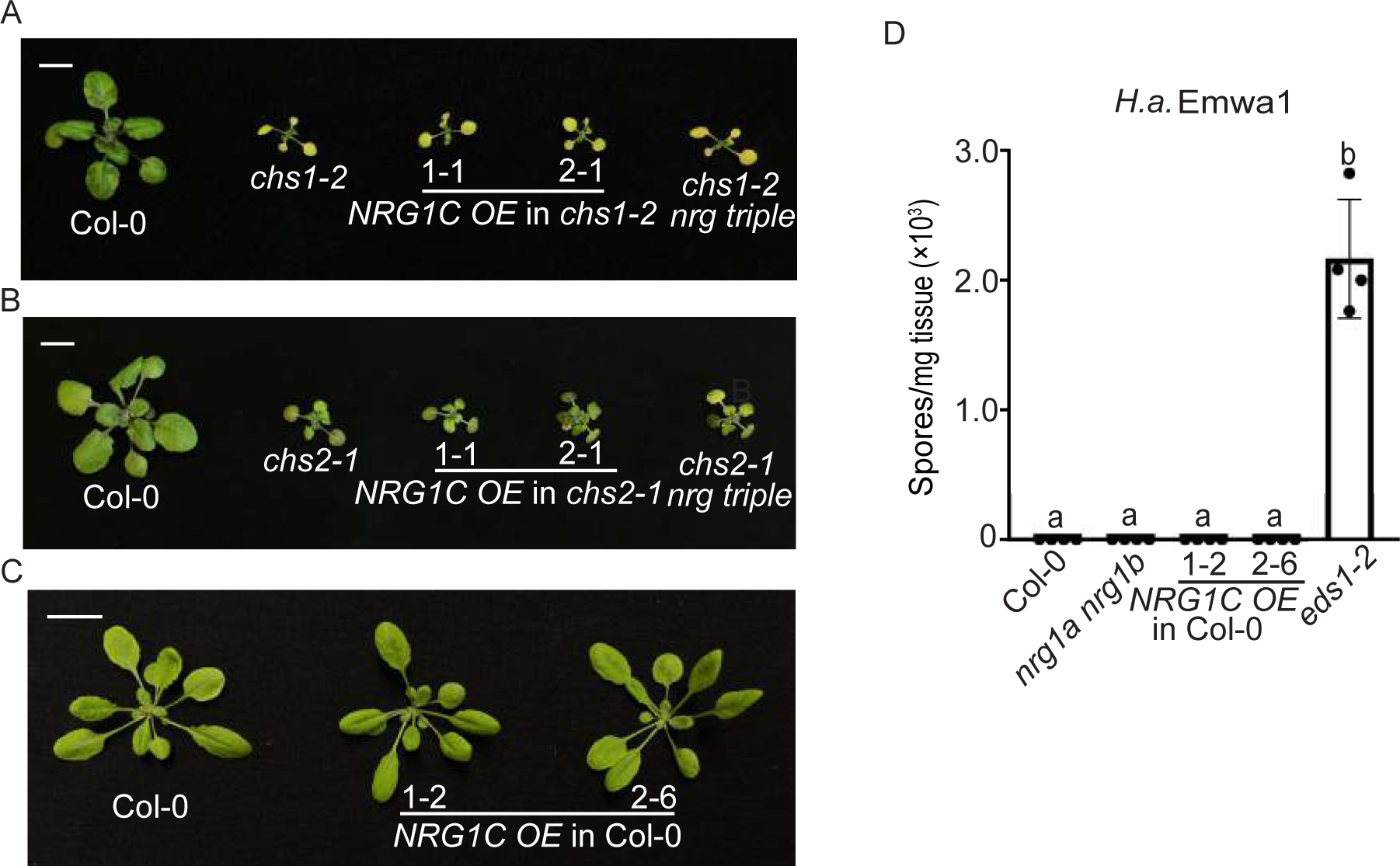
Overexpression of *NRG1C* has no effect on *chs1-2* or *chs2-1*-mediated autoimmunity. A-B. Morphology of four-week-old soil-grown plants of Col-0, *chs1-2* (A), *chs2-1* (B) and two independent transgenic lines of *NRG1C OE* into *chs1-2* (A) and *chs2-1* (B) backgrounds. Plants were grown at 16 °C under long day conditions. Bar = 1cm. (C) Morphology of four-week-old soil-grown plants of Col-0 and two independent transgenic lines of *NRG1C OE* in Col-0 background. Bar = 1cm. (D) Quantification of *H.a.* Emwa1 sporulation in the indicated genotypes at 7 dpi with 10^5^ spores per ml water. Statistical significance is indicated by different letters (*p* < 0.01). Error bars represent means ± SE (n=4). Three independent experiments were carried out with similar results.

**Supplemental Figure 6.**
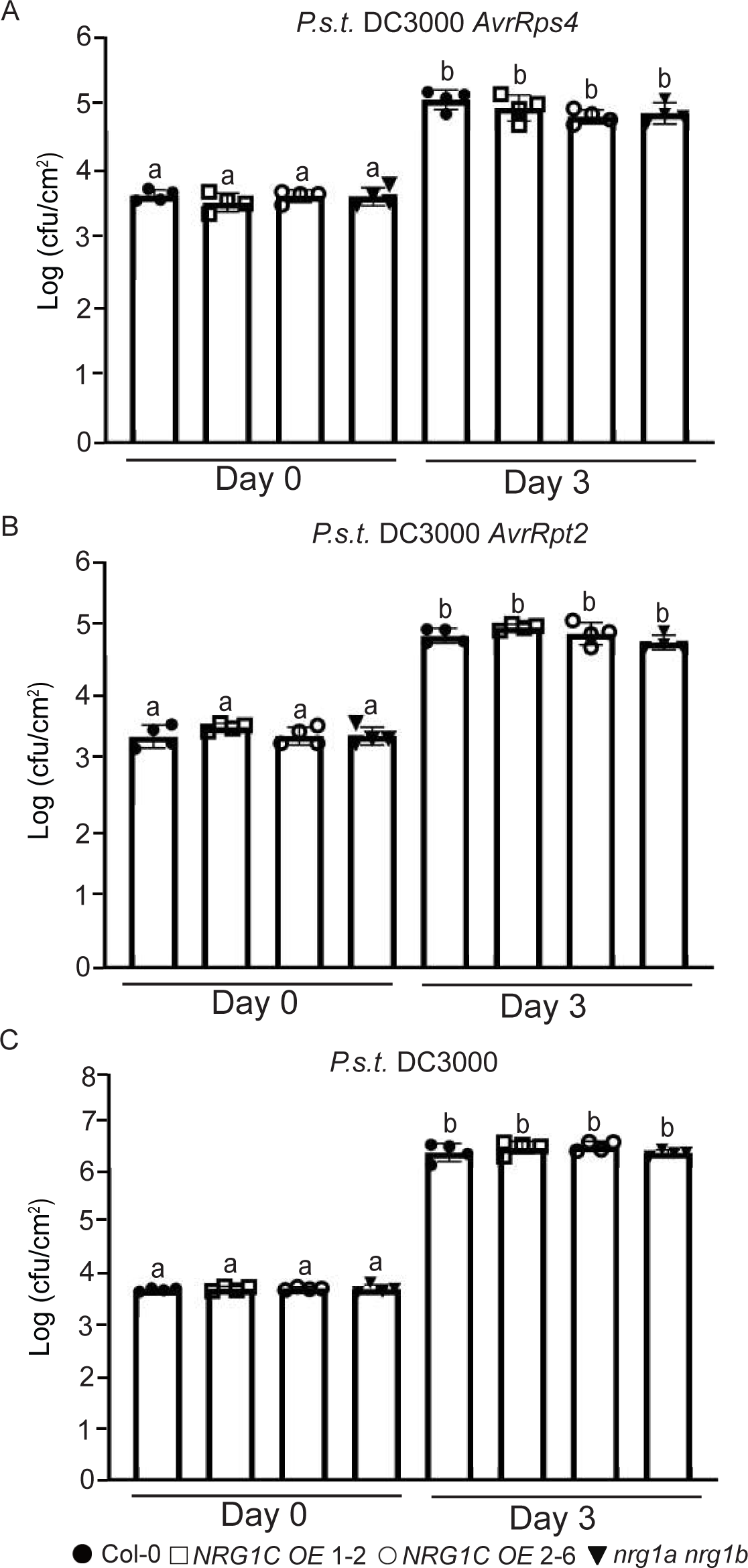
NRG1C does not affect TNL RPS4 or CNL RPS2-mediated bacterial growth and basal defense. A-C. Growth of *P.s.t. avrRps4* (A), *avrRpt2* (B) and *P.s.t.* DC3000 (C) in four-week-old leaves of the indicated genotypes at 0 dpi and 3 dpi with bacterial inoculum of OD600 = 0.001. Statistical significance is indicated by different letters (*p* < 0.01). Error bars represent means ± SE (n=4). Three independent experiments were carried out with similar results.

**Supplemental Figure 7.**
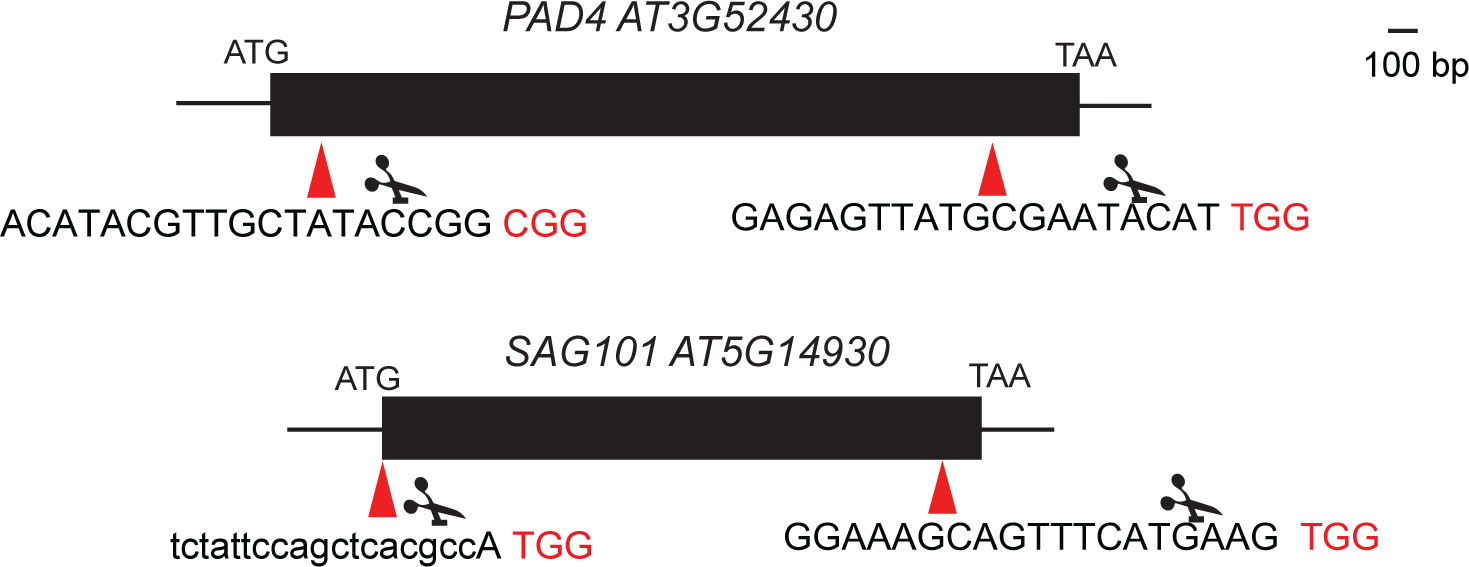
Schematic description of CRISPR/Cas9 construct design to delete *PAD4* **or *SAG101*.** Red arrows indicate the location of two guide RNA target sites. Target sequences are shown in the below. The letters in red indicate the NGG PAM sequences. Scissors indicate where the Cas9 cut the genomic sequence.

**Supplemental Figure 8.**
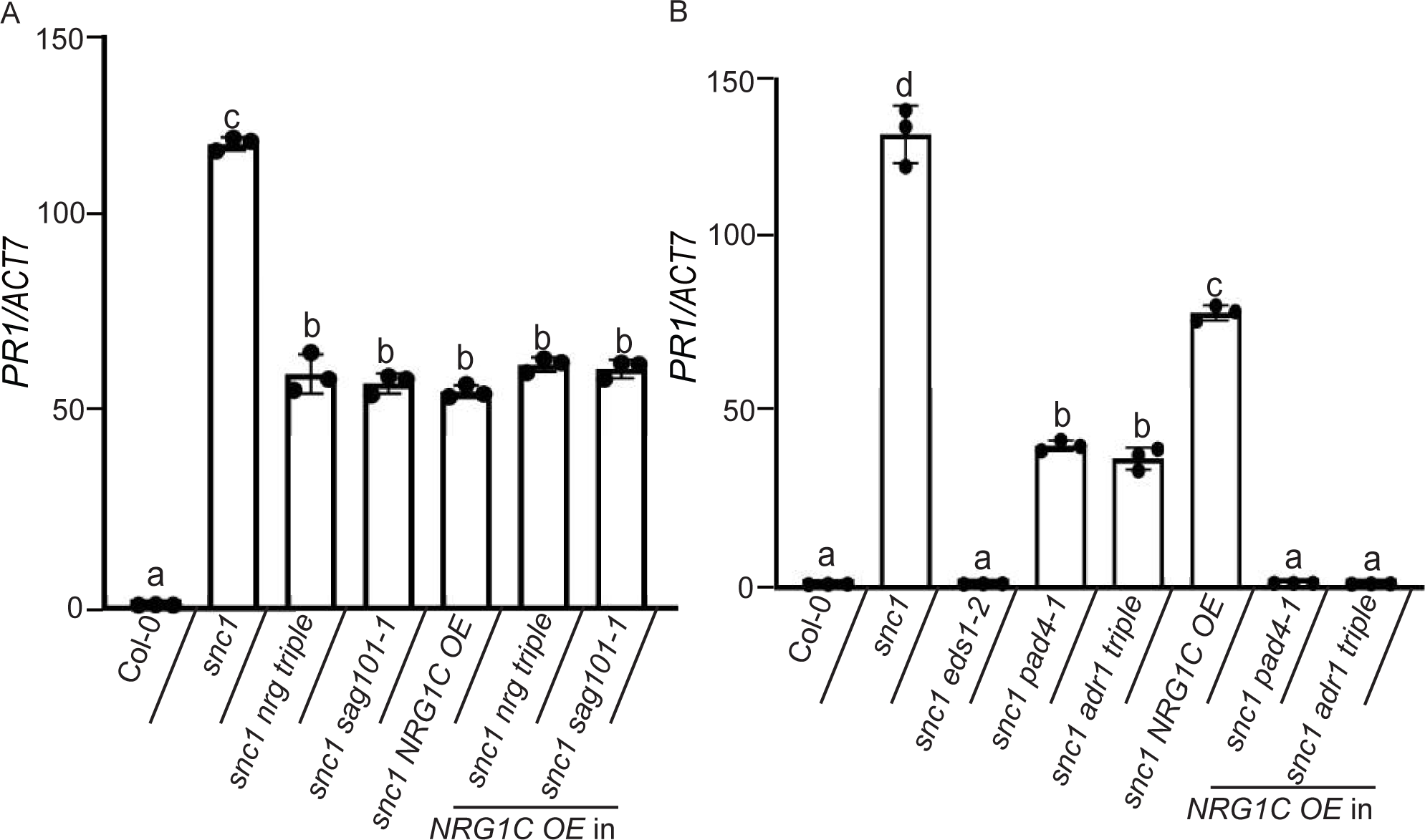
NRG1C works in the same pathway with the EDS1-NRG1-SAG101 module. A-B. *PR1* gene expression in the indicated genotypes as determined by RT-PCR (Col-0 serves as control whose *PR1* transcript level was set at 1.0). Statistical significance is indicated by different letters (*p* < 0.01). Error bars represent means ± SE (n = 3). Two independent experiments were carried out with similar results.

**Supplemental Figure 9.**
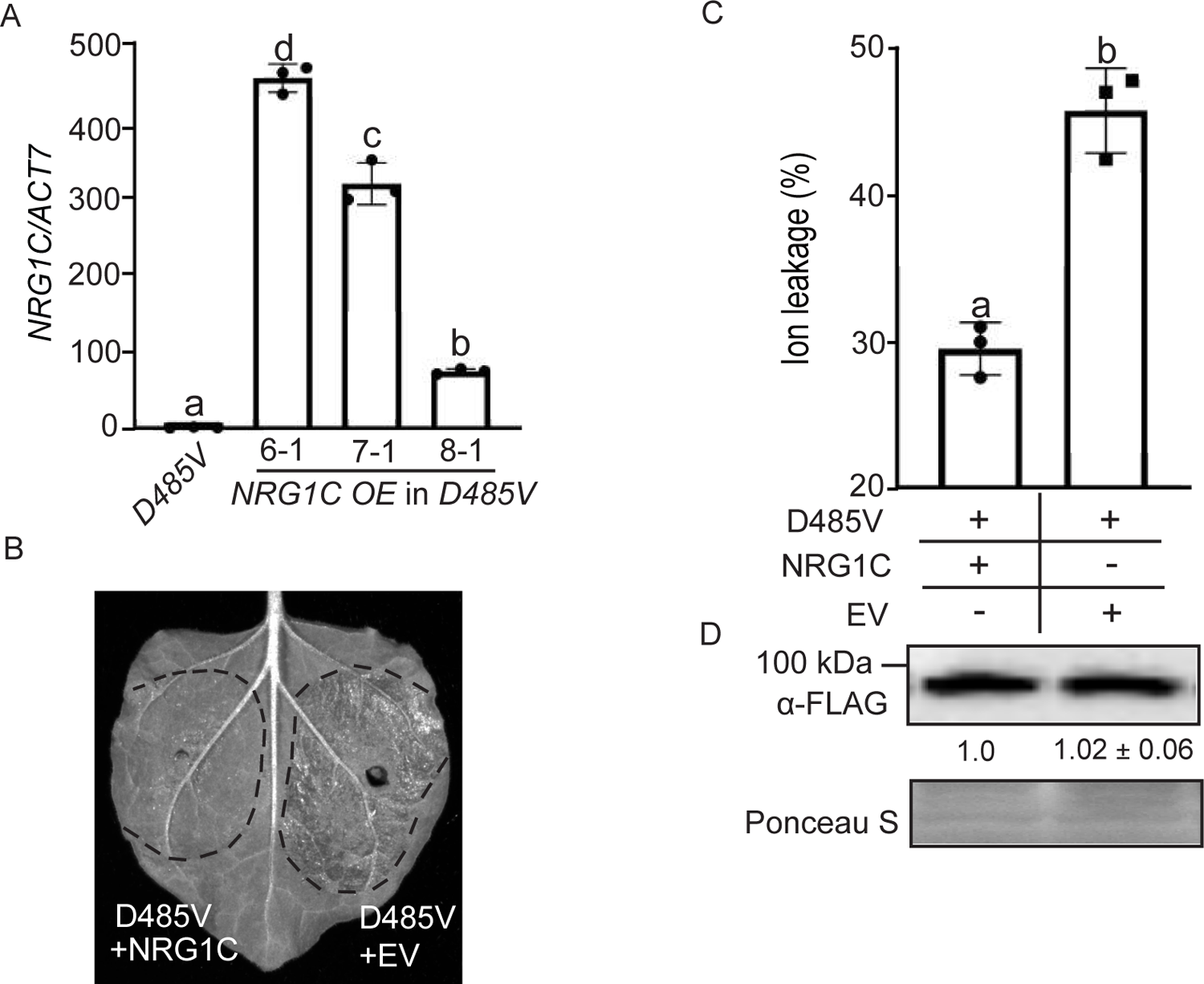
NRG1C antagonizes NRG1A D485V-mediated HR. (A) *NRG1C* gene expression in the indicated plants as determined by RT-PCR (*NRG1A D485V* serves as control whose *NRG1C* transcript level was set at 1.0). Statistical significance is indicated by different letters (*p* < 0.01). Error bars represent means ± SE (n = 3). Two independent experiments were carried out with similar results. (B) HR in the *N. benthamiana* leaves expressing NRG1A D485V-3FLAG and EV (Empty Vector) in the absence (right) or presence (left) of NRG1C. Photos were taken at 4 dpi. Three independent experiments were carried out with similar results. (C) Ion leakage of *N. benthamiana* leaves upon *Agrobacterium* infiltration under the same conditions as for B. Statistical significance is indicated by different letters (*p* < 0.01). Error bars represent means ± SE (n = 3). Three independent experiments were carried out with similar results. (D) Immunoblot analysis of protein levels of NRG1A D485V-3FLAG in leaves of B. Samples were harvest at 24 hpi, when no macroscopic HR was visible. Equal loading is shown by Ponceau S staining of a non-specific band. The numbers below represent the normalized ratio between the intensity of the protein band and the Ponceau S band ± SE (n=3). Molecular mass marker in kiloDaltons is indicated on the left.

**Supplemental Figure 10.**
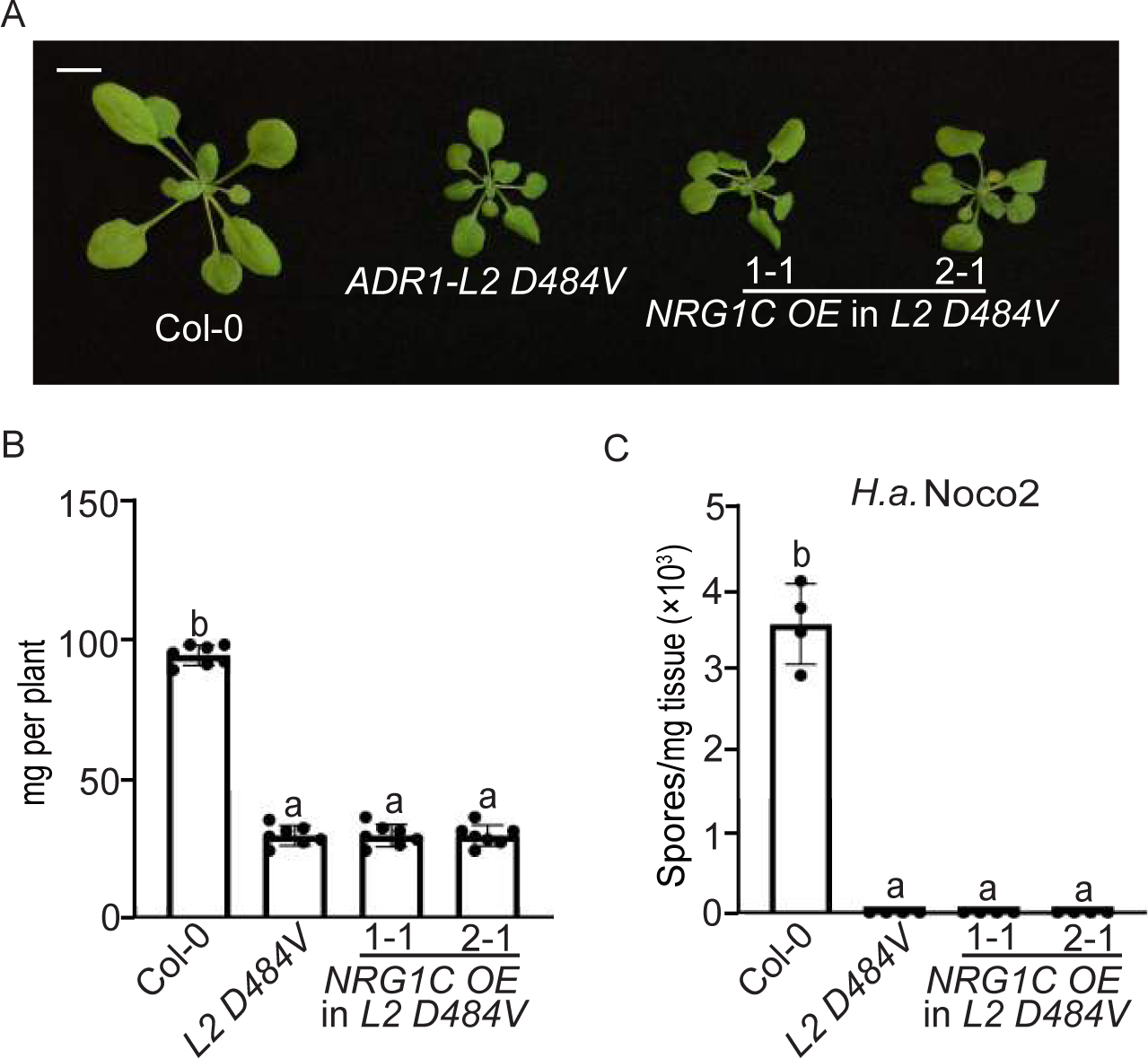
NRG1C does not influence the autoimmunity of *ADR1-L2 D484V*. (A) Morphology of three-week-old soil-grown plants of Col-0, *ADR1-L2 D484V*, and two independent transgenic lines of *NRG1C OE* into *ADR1-L2 D484V* background. *OE* stands for overexpression of *NRG1C*. Bar = 1cm. (B) Fresh weights of plants in (A). Statistical significance is indicated by different letters (*p* < 0.01). Error bars represent means ± SE (n=7). (C) Quantification of *H.a.* Noco2 sporulation in the indicated genotypes at 7 dpi with 10^5^ spores per ml water. Statistical significance is indicated by different letters (*p* < 0.01). Error bars represent means ± SE (n=4). Three independent experiments were carried out with similar results.

**Supplemental Figure 11.**
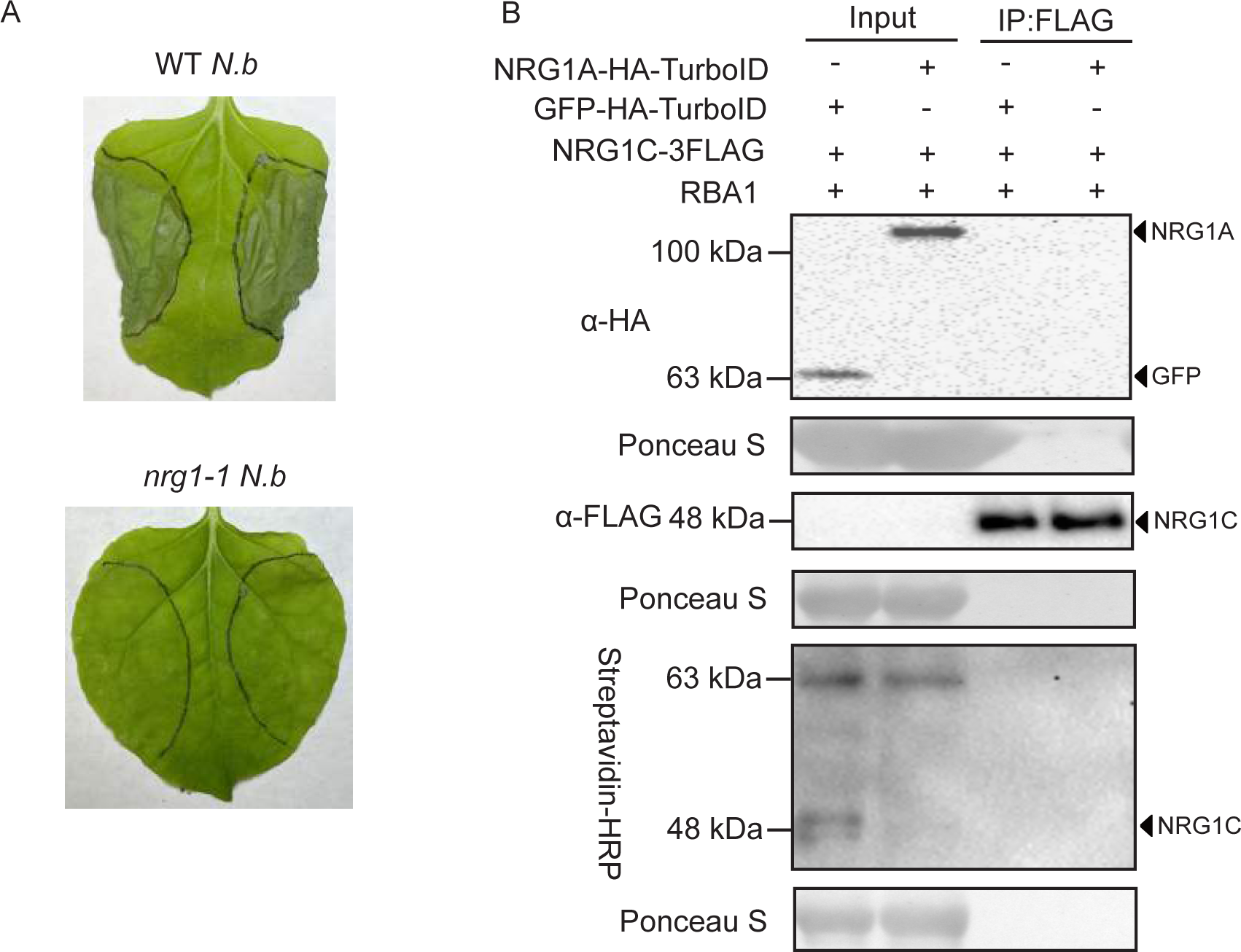
NRG1C does not associate with full-length NRG1A. (A) HR in the WT or *nrg1-1 N. benthamiana* leaves expressing RBA1. Photos were taken at 2 dpi. Three independent experiments were carried out with similar results. (B) Immunoprecipitation of NRG1C-3FLAG and biotinylation of NRG1C-3FLAG by NRG1A-HATurboID in *N. benthamiana* in the presence of RBA1. Immunoprecipitation was carried out with anti-FLAG beads. The 3FLAG-tagged proteins were detected using an anti-FLAG antibody. The HA-TurboID-tagged proteins were detected using an anti-HA antibody. The biotinylated proteins were detected using Streptavidin-HRP. Molecular mass marker in kiloDaltons is indicated on the left. Three independent experiments were carried out with similar results.

**Supplemental Figure 12.**
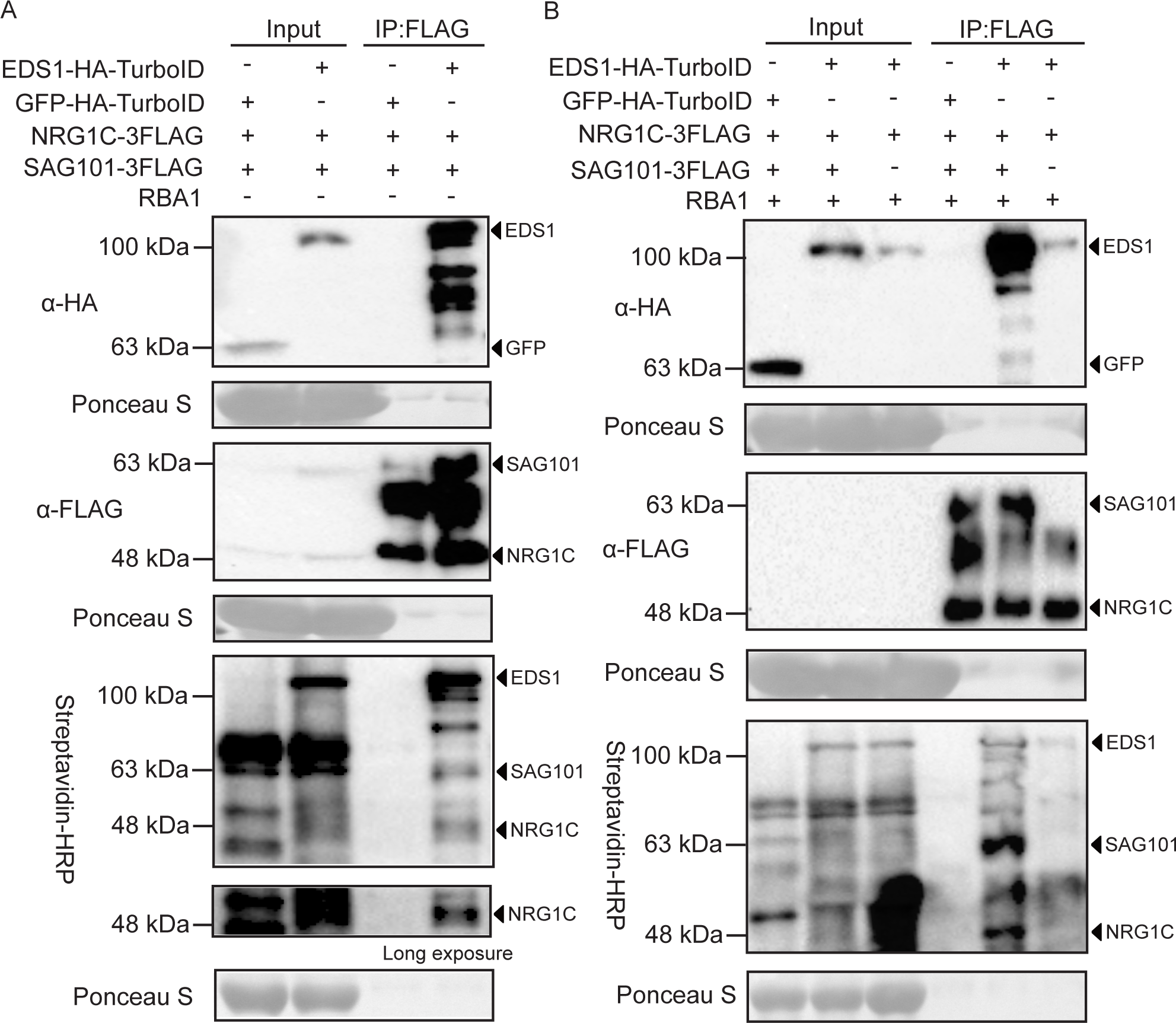
The association between NRG1C with EDS1-SAG101 complex depends on TIR induction and SAG101. (A) Immunoprecipitation and biotinylation of NRG1C-3FLAG and SAG101-3FLAG by EDS1-HA-TurboID in *N. benthamiana* in the absence of RBA1. Immunoprecipitation was carried out with anti-FLAG beads. The 3FLAG-tagged proteins were detected using an anti-FLAG antibody. The HA-TurboID-tagged proteins were detected using an anti-HA antibody. The biotinylated proteins were detected using Streptavidin-HRP. Molecular mass marker in kiloDaltons is indicated on the left. Three independent experiments were carried out with similar results. (B) Immunoprecipitation and biotinylation of NRG1C-3FLAG or/and SAG101-3FLAG by EDS1-HA-TurboID in *N. benthamiana*. Immunoprecipitation was carried out with anti-FLAG beads. The 3FLAG-tagged proteins were detected using an anti-FLAG antibody. The HA-TurboID-tagged proteins were detected using an anti-HA antibody. The biotinylated proteins were detected using Streptavidin-HRP. Molecular mass marker in kiloDaltons is indicated on the left. Three independent experiments were carried out with similar results.

**Supplemental Figure 13.**
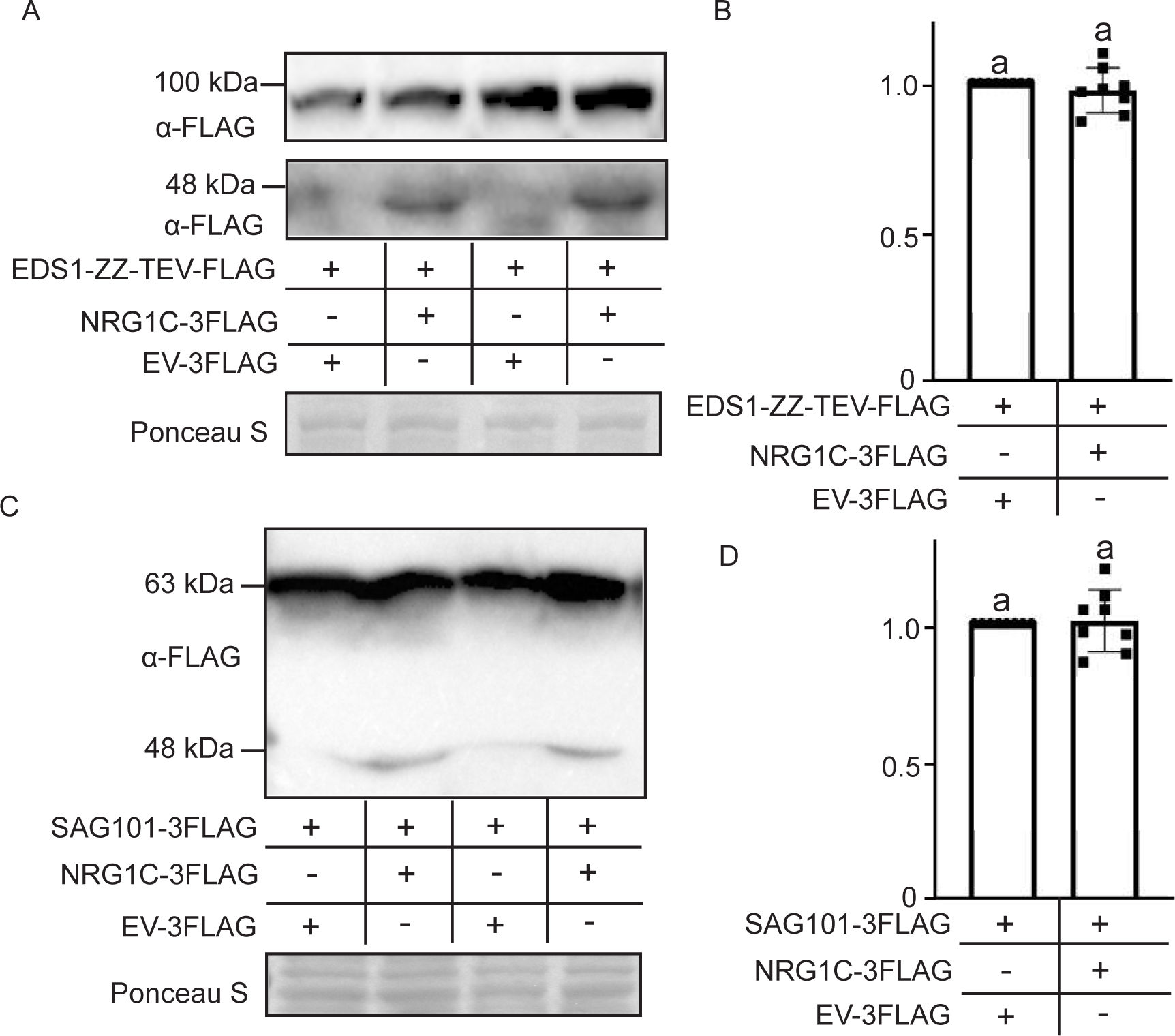
Overexpression of *NRG1C* does not affect the protein levels of EDS1 and SAG101. (A) Immunoblot analysis of protein levels of EDS1-ZZ-TEV-FLAG in the absence or presence of NRG1C-3FLAG. Equal loading is shown by Ponceau S staining of a non-specific band. Molecular mass marker in kiloDaltons is indicated on the left. (B) Quantification of band intensity of (A). The numbers represent the normalized ratio between the intensity of the protein band and the Ponceau S band ± SE (n=8). (C) Immunoblot analysis of protein levels of SAG101-3FLAG in the absence or presence of NRG1C-3FLAG. Equal loading is shown by Ponceau S staining of a non-specific band. Molecular mass marker in kiloDaltons is indicated on the left. (D) Quantification of band intensity of C. The numbers represent the normalized ratio between the intensity of the protein band and the Ponceau S band ± SE (n=8).

**Supplemental Figure 14.**
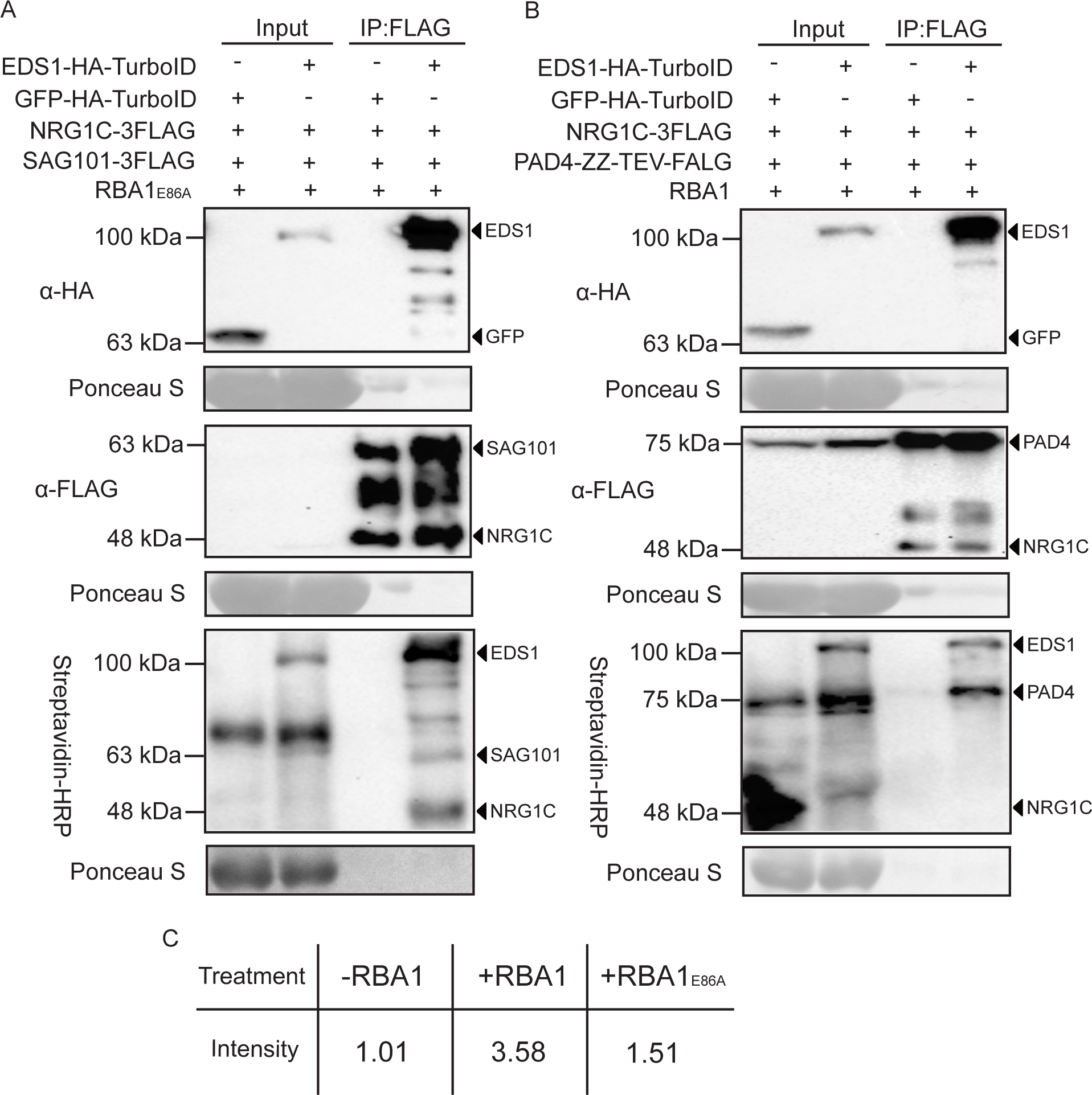
TIR NADase enzyme activity promotes the association between NRG1C with EDS1-SAG101 complex, and NRG1C does not associate with the EDS1-PAD4 complex. (A) Immunoprecipitation and biotinylation of NRG1C-3FLAG and SAG101-3FLAG by EDS1-HA-TurboID in *N. benthamiana* in the presence of RBA1E86A. Immunoprecipitation was carried out with anti-FLAG beads. The 3FLAG-tagged proteins were detected using an anti-FLAG antibody. The HA-TurboID-tagged proteins were detected using an anti-HA antibody. The biotinylated proteins were detected using Streptavidin-HRP. Molecular mass marker in kiloDaltons is indicated on the left. Three independent experiments were carried out with similar results. (B) Immunoprecipitation and biotinylation of NRG1C-3FLAG and PAD4-ZZ-TEV-FLAG by EDS1-HA-TurboID in *N. benthamiana* in the presence of RBA1. Immunoprecipitation was carried out with anti-FLAG beads. The ZZ-TEV-FLAG and 3FLAG-tagged proteins were detected using an anti-FLAG antibody. The HA-TurboID-tagged proteins were detected using an anti-HA antibody. The biotinylated proteins were detected using Streptavidin-HRP. Molecular mass marker in kiloDaltons is indicated on the left. Three independent experiments were carried out with similar results. (C) The normalized ratio between the intensity of the NRG1C band and the SAG101 band in different treatments. As EDS1 constitutively associates with SAG101, SAG101 bands were used as controls.

**Supplemental Figure 15.**
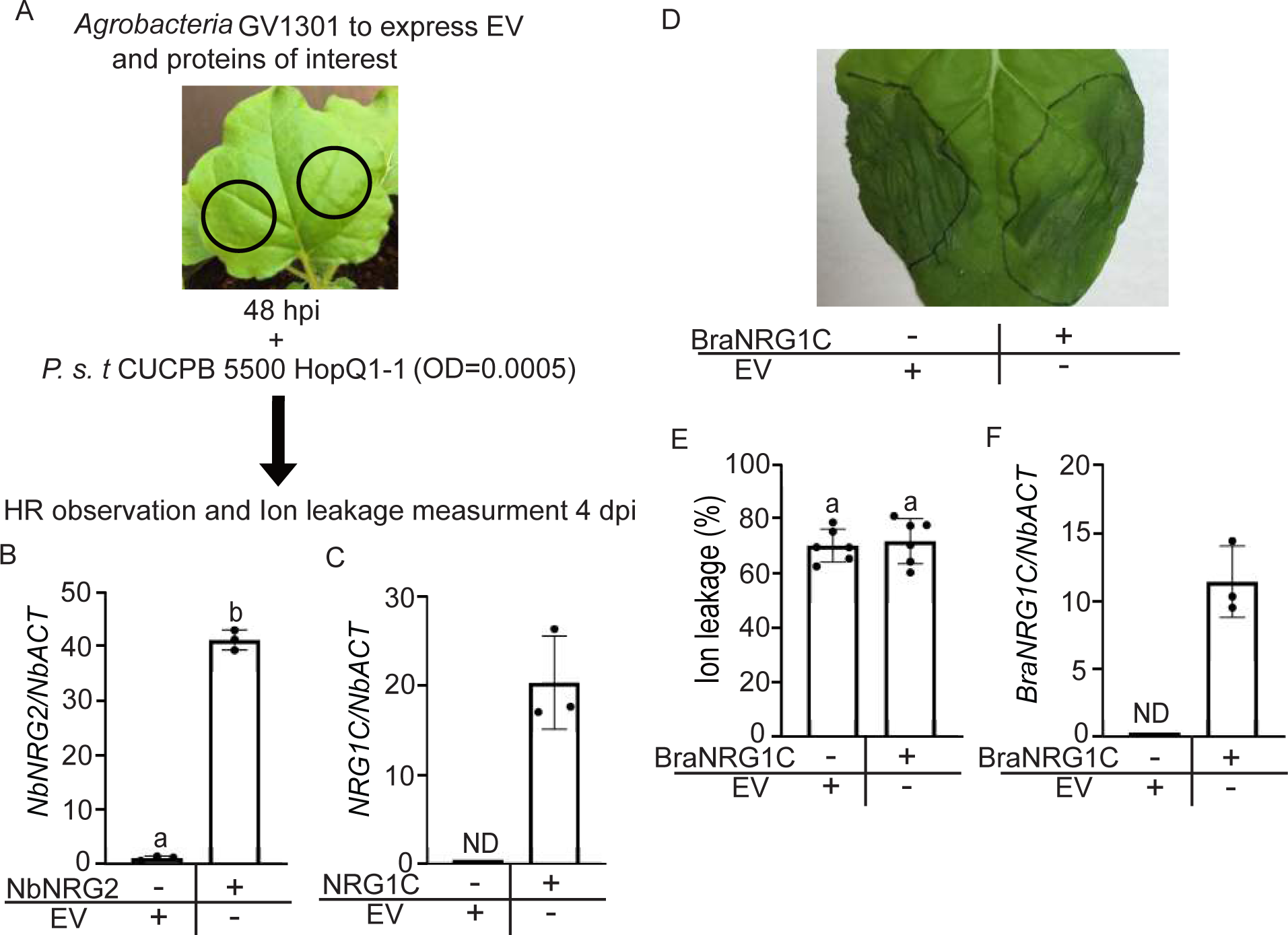
Brassicaceae NRG1C ortholog does not influence HopQ1-1-mediated HR in *N. benthamiana*. (A) Schematic showing *N. benthamiana* transient expression assays to test protein functionalities in HopQ1-1-dependent HR by infiltrating with *P.s.t.* CUCPB5500 HopQ1-1 (see Methods for details). B-C. *NbNRG2* (B) and *NRG1C* (C) genes expression in the indicated leaves as determined by RT-PCR. Statistical significance is indicated by different letters (*p* < 0.01). Error bars represent means ± SE (n = 3). The *NbNRG2* transcript level in the presence of EV treatment was set at 1.0. Two independent experiments were carried out with similar results. (D) HR in the *N. benthamiana* leaves expressing the indicated proteins at 48 hpi following the infiltration of *P.s.t.* CUCPB5500 HopQ1-1 to deliver HopQ1-1 effector. Photographs were taken at 4 dpi after *P.s.t.* CUCPB5500 HopQ1-1 infiltration. (E) Ion leakage of *N. benthamiana* leaves upon *P.s.t.* CUCPB5500 HopQ1-1 infiltration in the same conditions as for the photographs in D. Statistical significance is indicated by different letters (*p* < 0.01). Error bars represent means ± SE (n = 6). Three independent experiments were carried out with similar results. (F) *BraNRG1C* gene expression in the indicated leaves as determined by RT-PCR. Error bars represent means ± SE (n = 3). Two independent experiments were carried out with similar results.

**Supplemental Figure 16.**
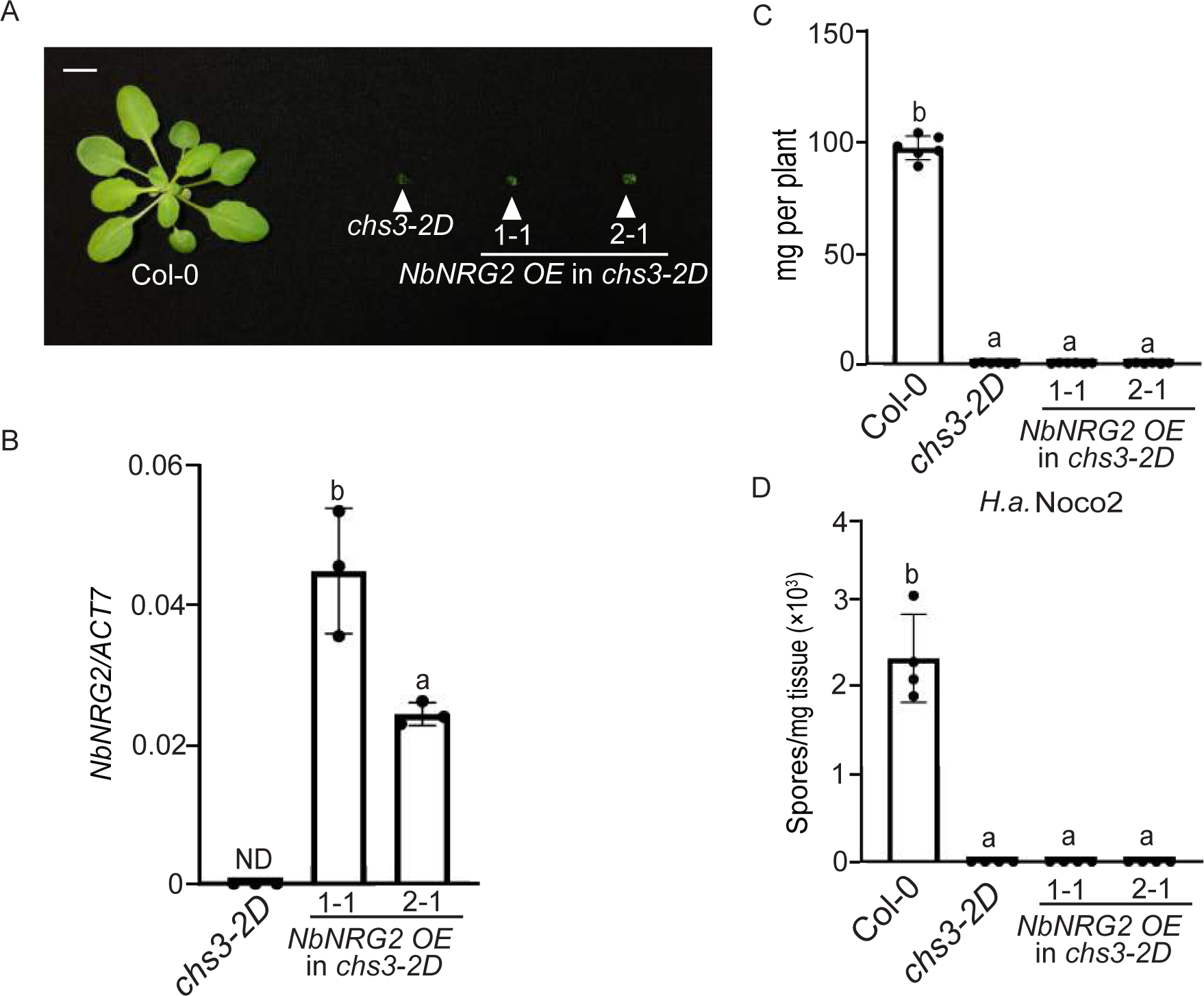
NbNRG2 does not influence *chs3-2D-*mediated autoimmunity. (A) Morphology of four-week-old soil-grown plants of Col-0, *chs3-2D*, and two independent transgenic lines of *NbNRG2 OE* into *chs3-2D* background. *OE* stands for overexpression of *NbNRG2*. Bar = 1cm. (B) *NbNRG2* expression level in the indicated plants as determined by RT-PCR. Statistical significance is indicated by different letters (*p* < 0.05). Error bars represent means ± SE (n = 3). Three independent experiments were carried out with similar results. (C) Fresh weights of plants in (A). Statistical significance is indicated by different letters (*p* < 0.01). Error bars represent means ± SE (n=6). (D) Quantification of *H.a.* Noco2 sporulation in the indicated genotypes at 7 dpi with 10^5^ spores per ml water. Statistical significance is indicated by different letters (*p* < 0.01). Error bars represent means ± SE (n=4). Three independent experiments were carried out with similar results.

**Supplemental Table 1.**
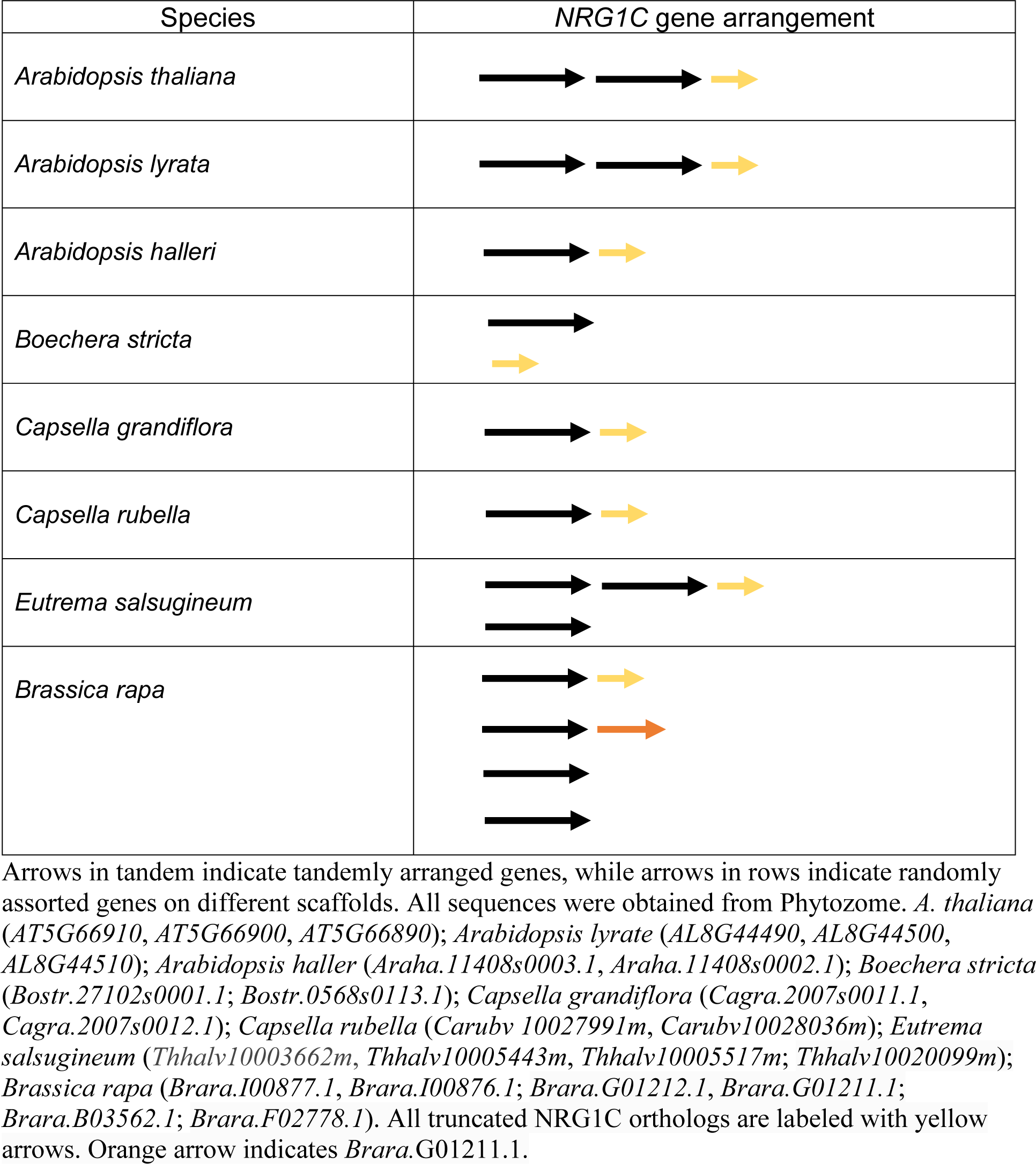
The genome arrangement of NRG1C in plant species.

**Supplemental Table 2.**
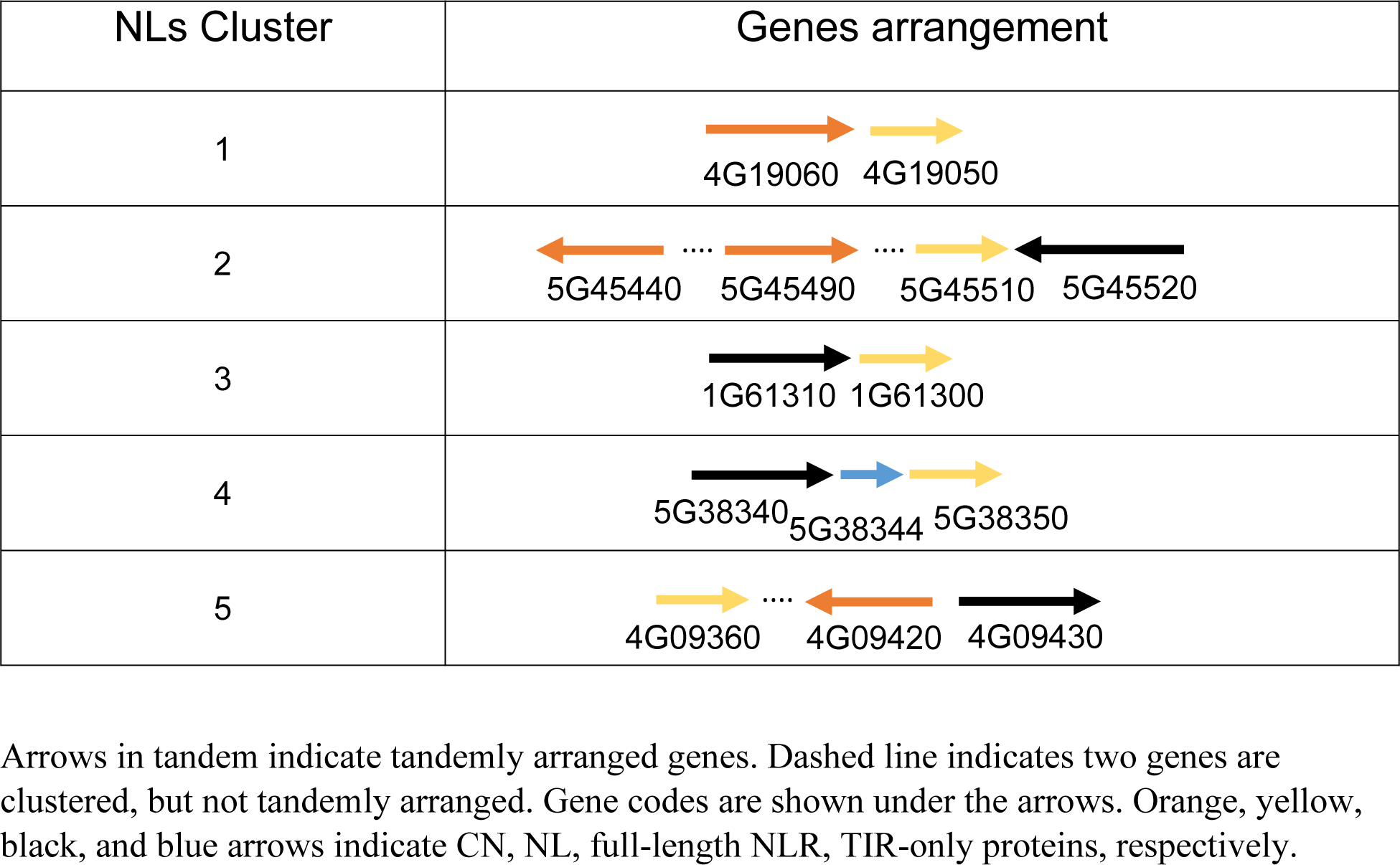
The genome arrangement of *NL* truncated genes in *A. thaliana*.

| NLs Cluster | Genes arrangement |
| --- | --- |
| 1 | 4G19060 4G19050 |
| 2 | 5G45440 5G45490 5G45510 5G45520 |
| 3 | 1G61310 1G61300 |
| 4 | 5G38340 5G38344 5G38350 |
| 5 | 4G09360 4G09420 4G09430 |

Arrows in tandem indicate tandemly arranged genes. Dashed line indicates two genes are clustered, but not tandemly arranged. Gene codes are shown under the arrows. Orange, yellow, black, and blue arrows indicate CN, NL, full-length NLR, TIR-only proteins, respectively.

**Supplemental Table 3.**
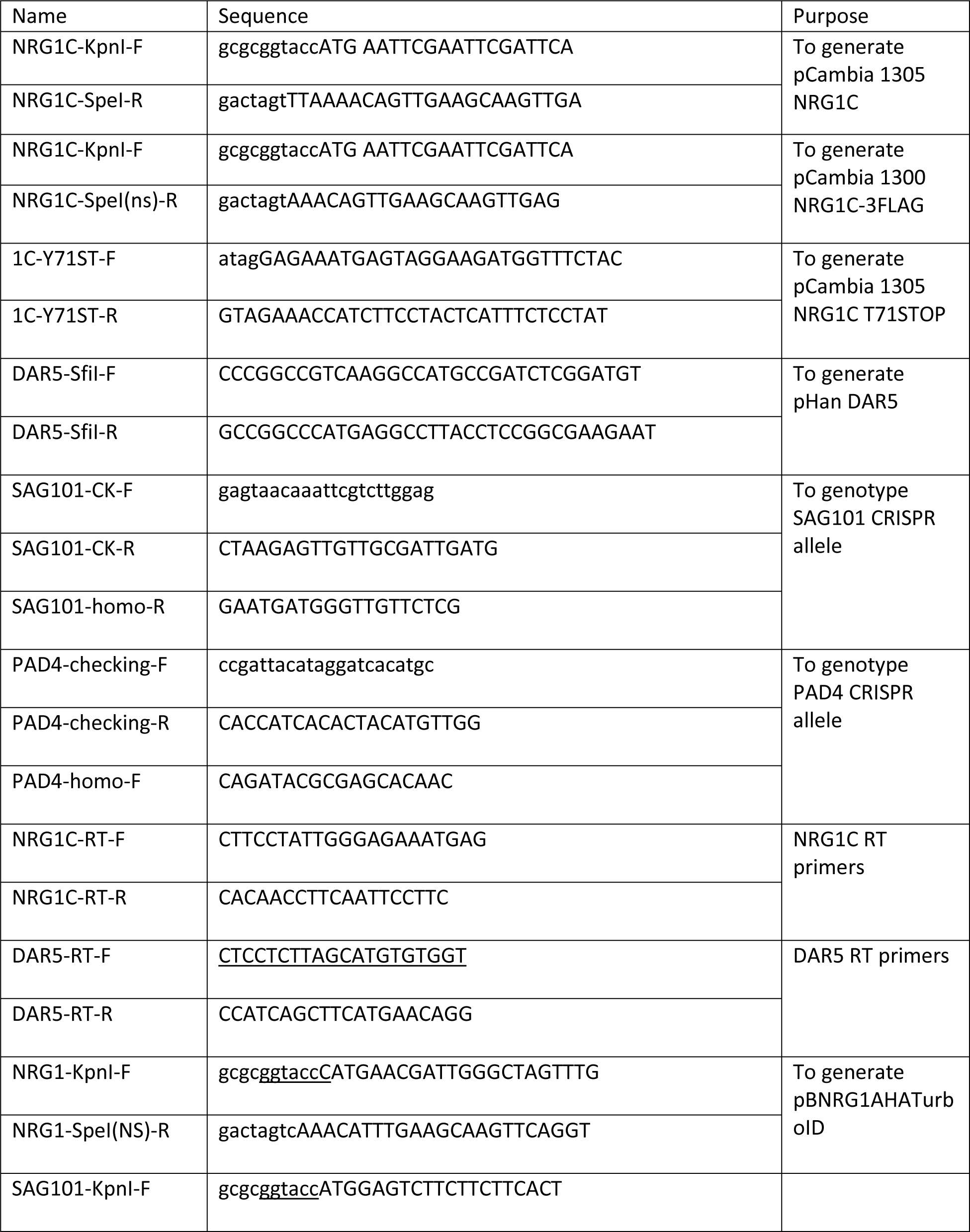
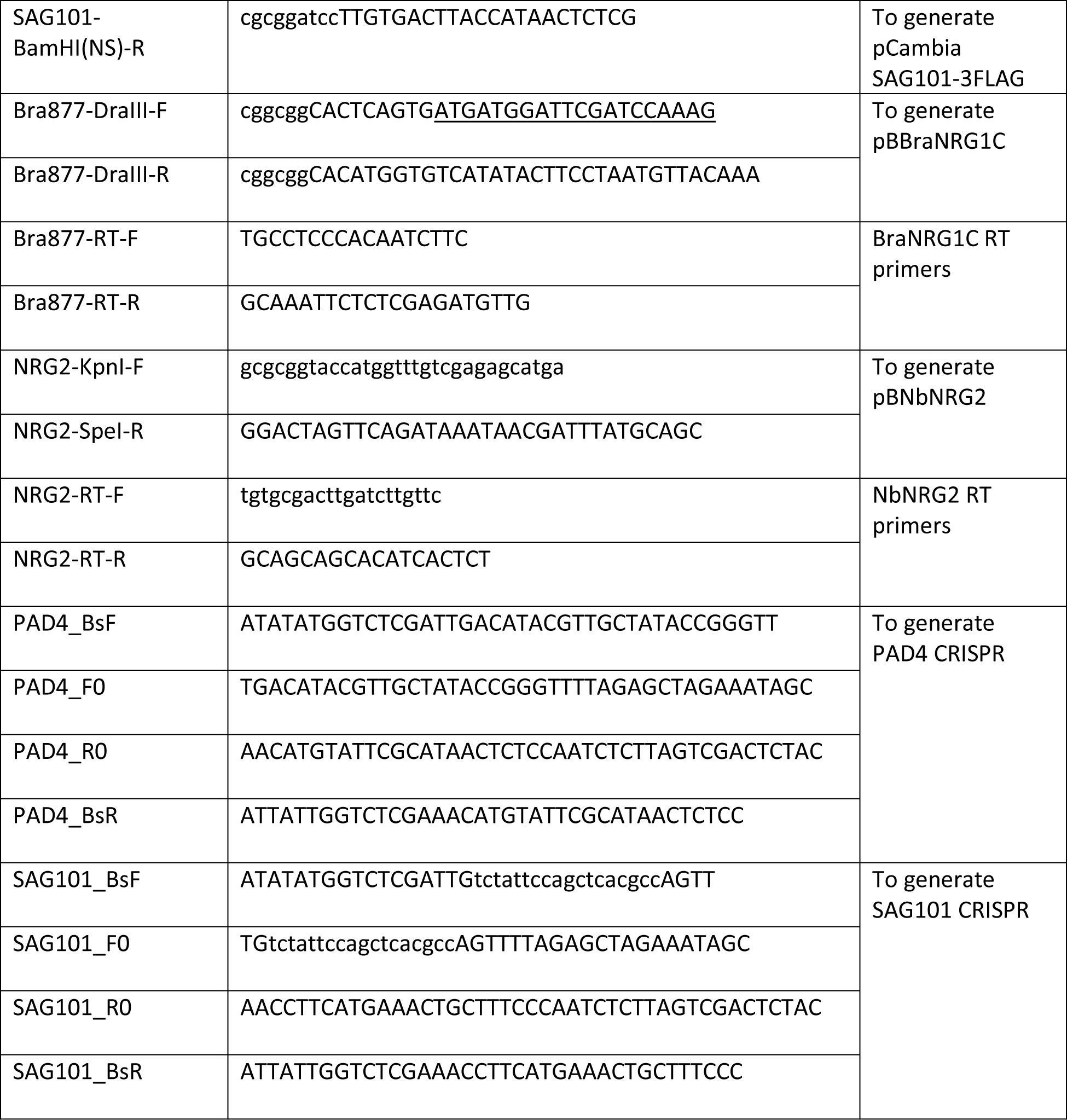
The list of primers used in this study.

